# Low salinity destabilizes the bacterial community of sugar kelp, *Saccharina latissima* (Phaeophyceae)

**DOI:** 10.1101/2023.12.07.570704

**Authors:** Siobhan Schenk, Connor Glen Wardrop, Laura Wegener Parfrey

## Abstract

As climate change progresses, the intensity and variability of freshwater outflow into the ocean are predicted to increase. The resulting increase in low-salinity events will be a source of stress for *Saccharina latissima* and potentially *Saccharina*-associated bacteria. Bacteria influence host health and can facilitate or hinder host survival and acclimation to stressful abiotic conditions. Therefore, understanding how bacterial communities change under abiotic stress is critical to understand how abiotic stress will affect kelp physiology. We investigated the effect of low-salinity stress on *Saccharina*-associated bacteria and the host by surveying the bacterial community associated with *Saccharina* and the surrounding environment across naturally occurring salinity gradients during the spring freshet across two years at four field sites with contrasting salinity profiles around Vancouver, Canada (519 samples), coupled with salinity manipulation experiments repeated eight times (269 samples). Overall, *Saccharina* harbours a stable core bacterial community, which decreases in relative abundance under abiotic stress. In the field, both salinity and temperature shape the bacterial community, with temperature having higher explanatory power most of the time. In the lab, we confirm that the patterns observed in the field can be replicated by manipulating salinity alone. Decreased relative abundance of core bacteria and increased community dissimilarity in low salinity, in both the lab and field, suggest that host filtering is significantly impaired in low salinity. In the context of a stable host-associated core bacterial community during non-stressful conditions, the change in the community composition observed during conditions of abiotic stress indicates a host stress response.

## Introduction

Kelp (*Laminariales*, brown algae) are marine foundation species that form dense underwater forests across the globe, providing food and habitat for other organisms (Steneck et al., 2002). Through ecosystem services and commercial harvest, kelp contribute an estimated $674 billion CAD per year to the global economy (Eger et al., 2023). However, kelp are threatened by global change factors, such as increased frequency and intensity of severe weather events, higher ocean temperatures, lower pH (Bindoff et al., 2019), increased coastal water turbidity (Filbee-Dexter et al., 2019), and lower ocean salinity (Davis et al., 2022; Filbee-Dexter et al., 2019; Andersen et al., 2011). All of these are associated with reductions in kelp recruitment, reproduction, and survival. Thus, kelp and the ecosystem services they provide are vulnerable to climate change (Bindoff et al., 2019; Filbee-Dexter et al., 2019; Goldsmit et al., 2021).

There is a large body of evidence demonstrating that low salinity is stressful to kelp. Field observations from Australia (Davis et al., 2022) and Norway (Andersen et al., 2011) find up to 100% mortality of kelp sporophytes after freshwater floods, followed by a re-growth of kelp in the following months. Lab studies also show that low salinity is stressful for kelp. Research of the microscopic gametophyte stage finds lower spore settlement and germination rates in low salinity (Lind & Konar, 2017). For the macroscopic sporophyte stage, lower photosynthetic efficiency (Bollen et al., 2016; Karsten, 2007), nitrogen uptake rates (Kumar et al., 2018), and growth rates (Mansilla et al., 2014) are reported in low-salinity treatments. As anthropogenic climate change progresses, lower ocean salinity has been predicted by modelling studies to be one of the most intense (Bindoff et al., 2019; Filbee-Dexter et al., 2019; Goldsmit et al., 2021) and least predictable abiotic stressors that kelp will experience (Bindoff et al., 2019).

The bacterial community of macroalgae can influence growth and morphology (Marshall et al., 2006; Provasoli & Pintner, 1980) and tolerance of abiotic conditions (Dittami et al., 2016). One unknown is how changes in abiotic stressors, including low salinity, will influence the kelp-associated bacterial community, and whether these changes exacerbate or ameliorate kelp stress tolerance.

Salinity is a strong determinant of bacterial community composition. In fact, a meta-analysis shows that host-associated bacterial community composition is primarily shaped by host association and salinity, even when other variables, including pH and temperature are taken into account (Lozupone & Knight, 2007). Field studies (van der Loos et al., 2023) and manipulative lab experiments (Saha et al., 2020; Stratil et al., 2014) show that salinity is an important factor in shaping the bacterial communities on non-kelp algal hosts. Studies of kelp (Lemay et al., 2018; Weigel & Pfister, 2019; Davis et al., 2023) and *Fucus* (Davis, 2022) have shown that seasonal changes and site differences are also important factors in shaping the bacterial community of brown algae. Time series in the field across multiple sites are necessary to disentangle these factors, particularly when paired with lab experiments to isolate the relative importance of various abiotic factors in altering the bacterial community (Trevathan-Tackett et al., 2019). The need for paired lab and field studies is apparent in the kelp literature. Field studies across sites at one time-point show that salinity, but not temperature significantly, alter the bacterial community associated with the kelp *Nereocystis* (Weigel & Pfister, 2019). However, in the lab, high temperatures alter the bacterial community of the kelp *Ecklonia* (Vadillo Gonzalez et al., 2024). We are not aware of lab-based studies that have examined how salinity alters the bacterial community of kelp.

Bacterial community composition typically changes in response to change in abiotic gradients across host-microbe systems, though the nature of these changes and whether they are correlated with positive, negative, or neutral host outcomes varies across systems. In studies examining the bacterial community of hosts under non-stressful salinity gradients, including in green algae (*Ulva sp.*) from localities with different salinity (van der Loos et al., 2023) and transplanted seagrasses (Adamczyk et al., 2022), changes in the bacterial community composition are not associated with changes in host condition. However, under stressful salinity gradients, changes in bacterial community composition may influence host functioning. For a freshwater strain of the brown alga *Ectocarpus sp.,* the bacterial community associated with the strain is essential to the alga’s ability to grow in fresh water (Dittami et al., 2016), showing that the bacterial community can improve brown algae’s ability to tolerate low salinity.

An important step in assessing the relationship between abiotic conditions and the bacterial community is to establish if the bacterial community is generally stable under the normal range of conditions, and only exhibits major changes under stressful conditions. A stable host-associated bacterial community is a requirement for the Anna Karenina Principle (AKP) to potentially apply. AKP predicts that in stressful conditions, beta diversity will increase (Zaneveld et al., 2017). Presumably, this destabilization is a result of the disruption of host filtering—host traits or mechanisms that selectively influence microbiome assembly—that typically maintain a stable, low-diversity bacterial community. Therefore, we predict that destabilization will also be accompanied by an increase in alpha diversity and a decrease in the relative abundance of core taxa. These patterns have been observed repeatedly in marine systems on corals (McDevitt-Irwin et al., 2017) and occasionally on sponges (Pita et al., 2018) in response to abiotic stress. However, a meta-analysis examining how temperature affects the bacterial community on a very wide range of taxa (aquatic and terrestrial) reveals that dissimilarity shows no significant change over temperature treatments and alpha diversity is more likely to decrease (Li et al., 2022), suggesting that AKP patterns and destabilization may not be the predominant stress response in the host-associated bacterial community. Alternatively, there may be a directional change in the bacterial community in stressful conditions. Establishing causality in the link between host condition and change in host-associated bacterial communities in response to stress is an open challenge that requires experimental manipulation (Egan et al., 2013; McDevitt-Irwin et al., 2017; Pita et al., 2018).

Field reciprocal transplant studies of corals across reef pools with different thermal profiles and a paired lab experiment (Ziegler et al., 2017) find long-term directional changes in the bacterial community (field) that are better adapted to the temperature profiles of the environment experienced by the host and its bacteria (lab). Lab studies of anemone bacterial communities (Baldassarre et al., 2022) find a similar pattern, where warm adapted corals are more resistant to heat stress than non-warm-adapted corals, mediated by the bacterial community, which stabilizes in the lab after 84 weeks of acclimation. Both studies control for the effect of host genetics and show that host-associated bacterial communities can show a directional change in the bacterial community towards an alternative stable state.

We conducted a two-year field survey from April to July in 2021 and 2022 and a concurrent lab study in 2022 to isolate the effect of salinity on the bacterial community the subcanopy kelp *Saccharina latissima* ((L.) C.E. Lane, C. Mayes, Druehl, and G. W. Saunders). Specifically, we tested if the observed shifts in the bacterial community are consistent with 1) a shift to a distinct low salinity community (turnover) or 2) destabilization of the community consistent with loss of host filtering. Existing data suggests that *Saccharina* has a relatively stable core bacterial community (King et al., 2022). If there is a shift to a distinct low-salinity community, we predict that the core bacterial community of *Saccharina* will be replaced by a new core bacterial community in stressful abiotic conditions. If the bacterial community is destabilized in low salinity, we predict an increase in community dissimilarity (beta diversity) in stressful abiotic conditions consistent with AKP principles. We also predict reduction of the core bacterial community and an increased alpha diversity, which in conjunction with increased beta diversity may indicate loss of host filtering in stressful conditions.

Our extensive dataset shows that the *Saccharina* bacterial community is largely stable with a consistent suite of core bacteria that are maintained across time, space, and salinity gradients. Layered on this broad pattern of stability, we find that salinity has a small but significant effect on the overall bacterial community composition in the lab and field, and that lower salinity significantly decreases the relative abundance of core bacteria, while increasing mean community dissimilarity. In the field, temperature generally has a stronger influence on the bacterial community than salinity. Together, these results suggest that host filtering by *Saccharina* is impaired in low salinity, and that high temperature and low salinity likely function as additive stressors.

## Methods

### Field site description

Five field sites near Vancouver, Canada, were visited across two successive years (Figure 1A). Both years, sites were sampled every two weeks (Figure 1B) at low tide, from mid-April until mid-July (2021) or early-August (2022) during the time of the annual freshwater influx caused by snow melt in late spring, hereafter called the spring freshet. We selected sites based on the presence of *Saccharina* and their salinity profiles through the freshet (Ryan et al., 2019), aiming for two sites that maintained relatively high salinity (above 20 psu) and two that dropped to salinities stressful for kelp (10 psu—15 psu); sites are numbered 1–5 from lowest to highest salinity. In 2021, we collected data at Site 1 (Lighthouse Park; 49.329°,-123.264°), Site 2 (Third Beach, Stanley Park; 49.302°,-123.158°); Site 3 (Sandy Cove Park; 49.333°,-123.222°), and Site 5 (Girl in Wetsuit, Stanley Park; 49.304°,-123.126°; Figure 1A). Salinity dropped below 15 psu at three of the four sites (Site 1, Site 2, and Site 3; Figure 1C), so in 2022, we replaced Site 1 with Site 4 (Cates Park; 49.300°,-122.958°; Figure 1A), which maintained higher salinity through the freshet and led to a more balanced sampling design. Within a sampling event, West Vancouver sites (Site 3 and Site 1 or Site 4) were always sampled on the same day and the Vancouver sites (Site 2 and Site 5) were sampled the subsequent day due to the tide height required and travel time.

**Figure 1.**
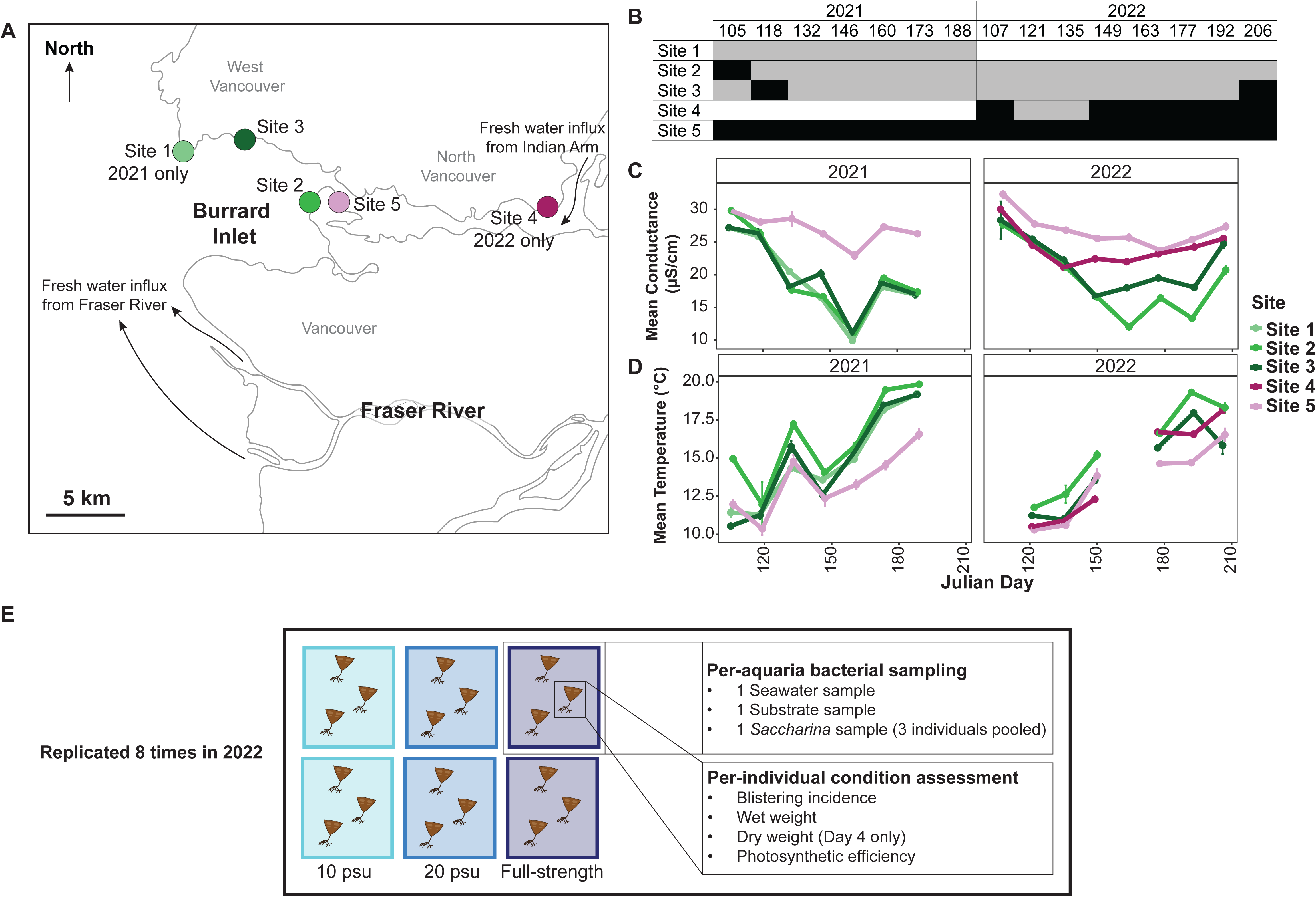
A) Map of the sampling sites with black arrows representing the path of main freshwater sources in the area. B) Table showing if a site was assigned as high (black) or low salinity (grey) for a field sampling round. The sampling rounds are numbered by the Julian Day of the first of the two sequential field days. White cells indicate that the site was not sampled. C) Conductivity (salinity) and D) water temperature profiles across sampling years with error bars showing +/-one standard deviation around the mean. E) Representative schematic of 2022 lab experiment. Note, in D, missing temperature data from 2022 are from days where a refractometer, rather than the YSI were used to measure water parameters. We include Julian day 206 in panels B, C, and D even though we exclude this sampling round in our field data analysis because experimental round 8 includes *Saccharina* collected during this sampling round. Also note, in E, the n = 1 for *Saccharina* bacterial samples because all individuals in the same aquaria are swabbed with the same cotton swab. Plots showing the correlation between salinity, temperature, and Julian Day are in Figure S1. Full temperature profiles from the temperature loggers show the same patterns across sites and time (Figure S2).

### Site conditions

In 2021 and 2022, we recorded water salinity (Figure 1C) and water temperature (Figure 1D) at the start of each site visit. In 2021, all measurements were conducted with an YSI ProQuatro Multiparameter Metre. On two sequential samplings in 2022, salinity only measurements were taken with a refractometer because the YSI was not available. In all cases, abiotic measurements were repeated three times per site visit and the measurement instrument was calibrated as per manufacturer’s instructions prior to each sampling.

### Temperature loggers and fake kelp installation

In 2022, two temperature loggers (ElectricBlue, EnvLogger with 27 mm, hardware v2.4) and two brown silicone *Saccharina* analogues (fake kelp hereafter, 30 cm long by 15 cm wide) per site were installed using an impact drill and underwater epoxy (GoToMarine, Z-Spar Splash Zone A-788 Kit). Temperature loggers were set at 0.5°C of precision and collected temperature data at a rate of every 30 minutes. One temperature logger was placed above and one below 1 m chart datum to capture the temperatures experienced by *Saccharina* across tide heights (Figure S2); the chart datum height is in the logger metadata. Both fake kelps at Site 4 and the higher intertidal fake kelps at Site 5 were lost partway through the freshet, likely removed by members of the public.

### Field bacterial data collection

In both years, two water samples, two rock swabs, and at least six *Saccharina* swabs were collected during each site visit to capture changes in the bacterial community throughout the spring freshet.

We swabbed the bottom 10 cm of the *Saccharina* thallus (the meristem region) because it is the newest and most selective tissue (Lemay et al., 2021), presumably hosting the bacterial community most indicative of *Saccharina.* Samples were taken wearing gloves sprayed with 70% ethanol between samples. Rocks were sampled to capture the background biofilm communities, for which we selected regions roughly 5 x 5 cm that were free of visible organisms. We gently rinsed the *Saccharina* and rocks with 0.22-μm filter-sterilized seawater before swabbing the surface for 10 s with a cotton-tipped swab (VWR, CA10805-154). The swab was then broken off into a cryovial (VWR, CA66021-993). After taking bacterial samples, we recorded the presence of any blisters on the kelp thallus. Blisters were observed only once on June 11, 2021, at Site 2 on three *Saccharina* individuals; the blistered tissue was not sampled.

Water column bacterial samples were collected by pre-filtering seawater through a 150-μm mesh before filtering the water through a 0.22-μm membrane (MilliporeSigma, Sterivex™ Filter Unit) until the filter clogged or 500 mL of water had been sampled. The Sterivex^TM^ was stored in a Whirl-Pak (VWR, 13500-390).

All samples were stored in a cooler with ice packs at −20°C until they could be brought to the lab (within three hours of sample collection), where they were then stored at −70°C until extraction.

### Saccharina field collection

The microbial communities of six *Saccharina* individuals were sampled by swabbing at each site visit. In 2021, individuals were selected haphazardly, with their position (out of the water, in the wave splash zone, fully submerged in the ocean, or in tide pools) recorded. These positions differ in conditions experienced, including desiccation, exposure, and in the case of tide pools, low water flow with a wider range of temperatures and oxygenation. All of these variables may influence the bacterial communities associated with *Saccharina.* We systematically tested the influence of being in a tide pool versus the ocean on the bacterial community of *Saccharina* in 2022 by sampling an additional six *Saccharina* from tide pools at Site 2 and compared them to individuals submerged in the ocean at this site. We used Site 2 because Site 2 consistently has accessible *Saccharina* in tide pools and the ocean, whereas the other sites do not. We used marginal PERMANOVAs including water salinity, water temperature, and position to test the influence of position in the intertidal on *Saccharina* bacterial communities. Neither position in the intertidal, in general (2021 position: pseudo-F_2,113_=1.071, R^2^=0.018 p=0.348), nor position in or out of tide pools (Site 2 in 2022 in or out of tide pool: QIAGEN pseudo-F_1,19_=1.5199, R^2^=0.059, p=0.160, Zymo pseudo-F_1,15_=0.944, R^2^=0.051, p=0.509) explained any additional variation in the bacterial community of *Saccharina*. Therefore, we have included *Saccharina* from all positions in our analyses. We only sampled *Saccharina* submerged in the ocean in 2022 at all other sites to increase consistency across samples.

### Lab experiment protocol

In 2022, we performed a manipulative lab experiment to isolate the effect of low-salinity stress (Figure 1E). We define stress as a condition that adversely affects growth via damage and/or resource allocation associated with damage prevention and cellular repair (Davison & Pearson, 1996; Harley et al., 2012). We performed eight experimental rounds through the 2022 field season, corresponding to each field-sampling event (Figure 1B), to assess the influence of salinity over time. After collecting the bacterial samples as described above, 18 *Saccharina* individuals were collected (the holdfast, stipe, and bottom 15 cm of the blade), numbered with flagging tape wrapped around the stipe, and transported in a cooler filled with seawater from Site 5 (approximately 1 h transit). *Saccharina* from Site 5 were used in the experiment because Site 5 has, by far, the largest population of *Saccharina,* and salinity stays relatively high.

*Saccharina* were incubated in seawater at 10°C on a 12 h light: 12 h dark photoperiod with bubblers to induce water motion. Each experimental round included six 8 L aquaria (two per treatment), with three *Saccharina* meristems per aquaria (Figure 1E), in one of three salinity treatments: 10 psu, 20 psu, and unaltered sea water (full-strength hereafter). The low-salinity treatment (10 psu) represents the lowest salinity observed at our sites (Figure 1C) and is reported to be stressful as measured by significantly lower effective quantum yield in sporophytes (Karsten, 2007). Significant increased death and damage were observed between 6 psu and 11 psu for *Saccharina* germlings (Peteiro & Sánchez, 2012). A salinity of 20 psu is commonly experienced at our sites (Figure 1C) and was previously associated with differential gene expression (Monteiro et al., 2019). The full-strength seawater is pumped from 30 m depth in Burrard Inlet and brought to UBC by truck. The salinity fluctuated between 31 psu and 32 psu depending on the experimental round (Table S1). Salinity was lowered by adding deionized water to the full-strength seawater as described previously (Gerard et al., 1987; Peteiro & Sánchez, 2012) and salinity was checked with a refractometer.

Most experimental rounds lasted four days with samples taken on Day 0 (in the field), Day 1 (24 h after collection), and Day 4. Our first experimental round lasted six days (Figure 2A). On Day 6, the *Saccharina* in 10 psu salinity died (turned green and fell apart to the touch; Figure 2A), so we shortened the incubation time to three days for experimental round two. We did not observe significant *Saccharina* damage on Day 3, so we extended the experimental duration to four days for the remaining experimental rounds (Figure 2A). For experimental round five, there are no dry weight measurements on Day 4 because we extended the experimental time to fourteen days, to observe whether the kelps in the 10 psu salinity would fall apart in as they did in round one. They did not.

**Figure 2.**
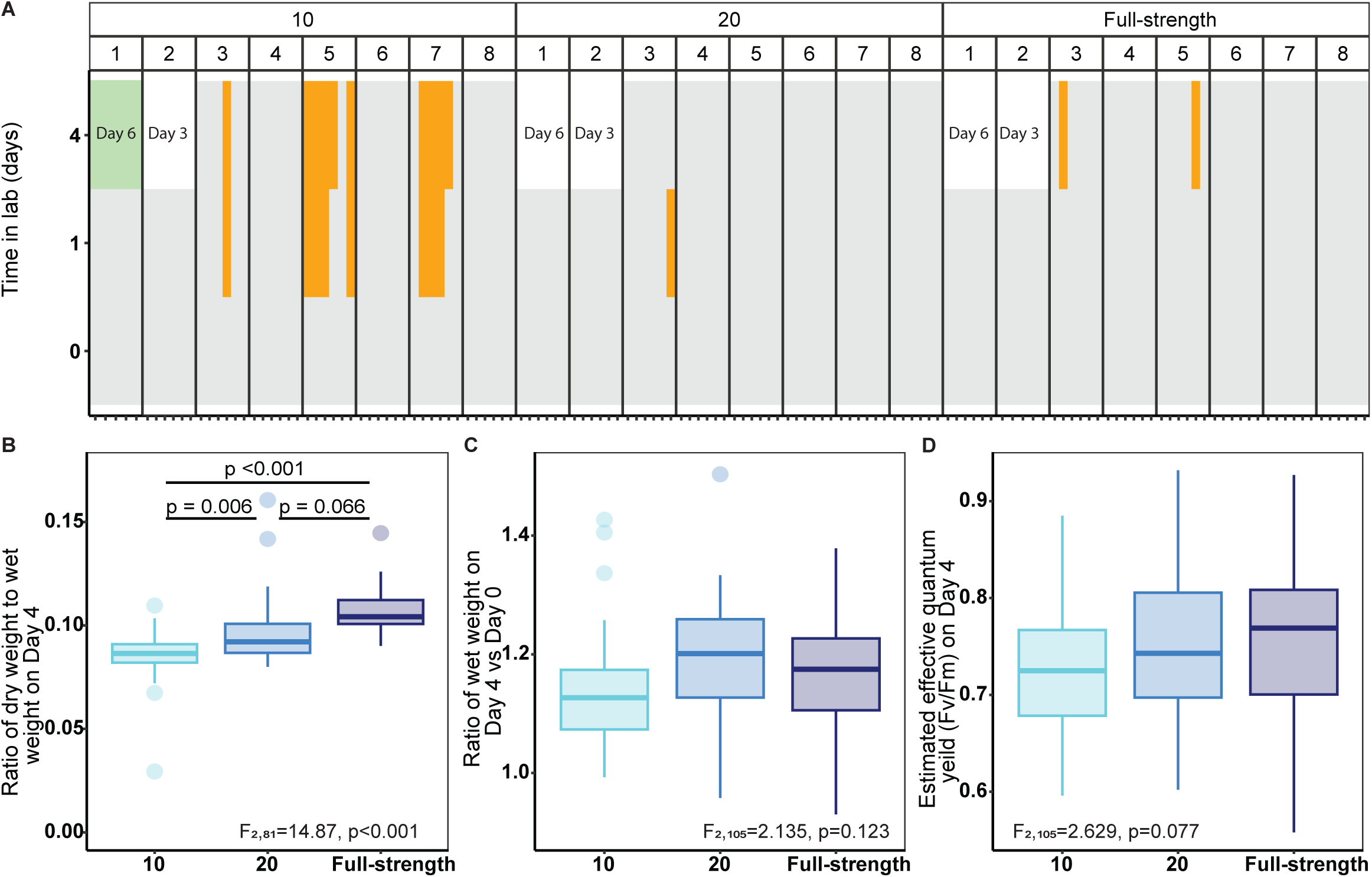
*Saccharina* stress phenotypes by lab salinity treatment. A) The presence (orange) or absence (grey) of blistering by lab experimental day, experimental round, and salinity treatment (in psu) for each *Saccharina* in lab. Experimental round 1 lasted six days and the *Saccharina* in the 10 psu condition became green and fell apart to the touch (indicated in green). Experimental round 2 lasted three days. Fisher’s Exact Test (p <0.001) followed by a Pairwise Fisher Test of the Day 4 data (experimental round 3 to 8), show that the 10 psu salinity has significantly more blistering than the 20 psu and full-strength salinity on Day 4 (p ≤ 0.036). B) The ratio of dry weight to wet weight on lab Day 4. C) The ratio of wet-weight on lab Day 4 to wet-weight on Day 0. D) The estimated effective quantum yield on lab Day 4 by salinity (in psu). For B, C, and D, ANOVA output comparing the different salinity treatments are indicated in the corresponding panel. Note, round 1 and 2 have no Day 4 observations for all panels. In addition, in panel B only, there are no Day 4 observations for experimental round 5 because we extended the experimental time to see if the *Saccharina* would turn green. After 14 days, green *Saccharina* was not observed in any salinity treatment. Day 1 comparisons for panels A, C, and D are in the supplementary results.

### Lab bacterial collection protocol

All lab bacterial sampling (Figure 1E) followed the protocol in the field section with the following modifications. The substrate samples were taken by swabbing the airline tubing of the bubbler rather than rocks and the aquaria water was not pre-filtered with a sieve before taking bacterial samples. Note the Day 0 *Saccharina* swabs are the Site 5 field samples. Since only six kelps are swabbed in the field, this means most lab experiment kelps are not swabbed on Day 0. Day 0 water and substrate samples were taken prior to adding the kelps to the aquaria. Lab *Saccharina* samples (Day 1 and Day 4) are swabs of all three kelps in the same aquaria to avoid pseudo-replication because we assumed the bacterial community was homogenized within the same aquarium (Chen & Parfrey, 2018). In total, we collected one of each sample type per aquaria, per lab sampling day (2 replicates for each treatment) and repeated the experiment eight times (Figure 1E).

### Lab Saccharina condition

After taking bacterial samples, the condition of each *Saccharina* was assessed on an individual basis (n=6 per salinity treatment/time point/experimental round) by quantifying blistering occurrence, wet weight, estimated quantum yield, and ratio of dry weight to wet weight on the last day of each experimental round (Figure 1E). These measurements serve as a phenotypic read-out to test whether the biologically relevant salinity treatments chosen for this experiment are stressful according to the definition presented above (Davison & Pearson, 1996; Harley et al., 2012) by testing the effect of salinity on growth in particular.

Blistering occurrence was recorded as in other kelp studies (Davis et al., 2022; Qiu et al., 2019). Estimated quantum yield measurements were taken with a Junior-PAM Chlorophyll Fluorometer as described in the user manual. Briefly, we calibrated the PAM and dark acclimated the *Saccharina* for five minutes on a damp cloth. Then, we placed the optical fibre over the thallus base (just above the stipe) and took the measurement. Wet weight measurements (holdfast, stipe, blade section) were taken after blotting dry the *Saccharina,* after which the kelp was promptly returned to its aquaria.

Dry weights were obtained as follows. After taking wet weight measurements on the last experimental day, stipes and holdfast were cut off from the blade section (Bollen et al., 2016). We weighed the blade sections before drying them in at 60°C for at least 24 h, and reweighed them (Bollen et al., 2016; van Ginneken, 2018).

### DNA extraction and sequencing

In 2021, all DNA was extracted with QIAGEN DNeasy PowerSoil Pro Kit 96-well plate; 47017 or single tube; 47014 following the manufacturer’s protocol. In 2022, 71% of field swabs (*Saccharina*, water, and rock) samples were extracted with QIAGEN 96-well kits. The remaining 29% of field samples were extracted with ZymoBIOMICS 96 MagBead DNA/RNA Kit (Zymo, R2136). All lab samples in 2022 were extracted with the Zymo kit. QIAGEN PowerSoil Max beads (QIAGEN, 12988-10) were used for the initial bead-beating step, and then the Zymo kit was used following the manufacturer’s protocol. The extraction method for each sample is indicated in the microbial metadata. The change was due to global supply chain issues. Each extraction plate included a blank swab or Sterivex^TM^ membrane (extraction blank, n=12).

Primers for PCR were 515F (5’-GTGYCAGCMGCCGCGGTAA-3’) and 806R (5’-GGACTACNVGGGTWTCTAAT-3’), which target the V4 region of the 16S rRNA gene. Both primers have an Illumina adaptor and golay barcode on the 5’ (Parada et al., 2016; Quince et al., 2011). PCR reactions were as follows: 1 μL extracted DNA (for extracted samples) or water (for PCR blanks, 1 per PCR plate, n=12), 15 μL Phusion™ High-Fidelity DNA Polymerase (Thermo Fisher, F530S), 2.4 μL of forward primer, 2.4 μL reverse primer (primers are from a 2.5 μM working stock solution), to a final reaction volume of 30 μL with molecular water (Thermo Fisher, SH30538FS). PCR reactions consisted of 30 s initial denaturation at 98°C, 25 cycles of amplification (30 s at 98°C, 30 s at 55°C, and 20 s at 72°C), and a final elongation step at 72°C for 10 min, which are the conditions recommended for the Phusion^TM^ polymerase. Samples where a band was not visible on a DNA gel after two attempts with 25 cycles were amplified with 35 cycles (indicated in the microbial metadata).

PCR reactions were cleaned with the QIAGEN QIAquick PCR Purification Kit (28104 or 28181), and quantification was performed with the Quan-IT PicoGreen assay kit (Thermo Fisher, P7589) following the manufacturer’s protocol. Samples were pooled to equal concentrations and sent for Bioanalyzer at the UBC Sequencing Facility, Vancouver, British Columbia. In 2021, the library was sent for Illumina MiSeq at the Hakai Institute, Campbell River. In 2022, both libraries were sent for illumina MiSeq at the UBC Sequencing Facility. In all cases, libraries were constructed with the Illumina, MS-1023003 kit (MiSeq v3, 2×300).

### Sequence data processing

Raw, demultiplexed reads were downloaded from the Illumina hub and imported into RStudio (R version 4.2.2, RStudio v2022.10.31; (Posit team, 2022; R Core Team, 2022)). Reads were processed following the dada2 pipeline (v1.24.0; Callahan et al., 2016). Primers were removed; reads were truncated to maintain high quality for downstream analysis with the filterAndTrim function (275 bp for forward and 200 bp for reverse in 2021 and 230 bp for forward and 175 bp for reverse in 2022); error rates were calculated; paired forward and reverse reads were merged; a sequence table was constructed; and chimeric reads were removed. The 2021 and 2022 sequence tables were merged (mergeSequenceTables) and amplicon sequence variants (ASVs) that differed by only the end base pair were collapsed (collapseNoMismatch), prior to assigning taxonomy with SILVA v138 (McLaren, 2020) formatted for the dada2 pipeline (McLaren & Callahan, 2021).

The merged sequence table, the taxonomy table, and metadata were grouped into a phyloseq object for filtering (v1.40.0; McMurdie & Holmes, 2013). Unassigned taxa and taxa assigned to chloroplasts, mitochondria, or eukaryotes were removed in addition to sequences assigned as *Pseudomonas*. *Pseudomonas* were present at high relative abundance in multiple extraction blanks and was abundant in nearly all samples extracted with the Zymo kit, but present in only 1 sample extracted with the QIAGENkits across both years; *Pseudomonas* is very likely a Zymo kit contaminant. Next, samples with fewer than 1000 reads were removed; then, ASVs representing less than 0.001% of total reads in the dataset were removed; counts in the ASV table that were five or less were converted to zero (per sample filtering, to minimize the effect of barcode switching); and ASVs found in less than two samples were removed. Finally, all samples from one 10 psu salinity aquarium in experimental round five were removed (four samples total) because we noticed that there was *Desmerestia viridis* lodged in the holdfast. *Desmerestia sp.* produce sulfuric acid (Eppley & Bovell, 1958) and pH alters the bacterial community of kelp (Qiu et al., 2019). We also removed the last field sampling event in 2022 (July 26^th^ and 27^th^) to make the time covered in 2021 and 2022 consistent as our sampling ended earlier in 2021. In total, filtering retained 17,438,672 of 24,862,382 total paired reads, 3,040 of an initial 29,105 ASVs, and 796 of the initial 1,095 samples, with a mean of 21,432 reads per sample. See table S2 for final sample number by year and site (field) or sample type (lab). We converted the data to an iNEXT (v3.0.0; Hsieh et al., 2022) compatible format with the metagMisc package (v0.0.4; Mikryukov, 2022) to perform coverage-based rarefaction. The sample coverage was set to 0.8 and iterated 1000 times.

We tested the influence of DNA extraction kit on diversity. By Kruskal-Wallis test comparing field *Saccharina*, rock, and water samples followed by a Benjamini-Hochberg correction for multiple comparisons, *Saccharina* (χ_1_ ^2^ = 120.07, p<0.001) and rock samples (χ ^2^ = 25.465, p<0.001) extracted with the Zymo kit had higher richness than those extracted with the QIAGENkit, but not water samples (χ ^2^ = 2.64, p=0.10). We tested for differences in beta diversity using PERMANOVA comparing samples extracted with Zymo versus QIAGEN for the same sample type and at a similar time of year (Julian Day 160 to 192; 3 sampling rounds). We found significant differences between extraction kits for *Saccharina*, rock, and water samples (Figure S3), indicating that the different extraction kits capture different bacterial communities. Therefore, we accounted for extraction kit in all analyses that include samples extracted with different kits (2022 field samples).

### Statistical analysis

To compare blistering incidence between salinity treatments, we used a Fisher’s exact test with a Benjamini-Hochberg correction with the package rstatix (v0.7.1; Kassambara, 2022b).

For all field bacterial data analysis, we include all 2021 sampling rounds (Julian day 105 to 188) and sampling rounds one to seven (Julian day 107 to 192). We exclude the sampling round starting on Julian day 206 from the field data analysis because it is not matched in the 2021 data. For all lab bacterial analysis, we include all 2022 rounds.

We ran permutational analyses of variance (PERMANOVA) on rarefied data with the adonis2 function in the package vegan (v2.6-4; Oksanen et al., 2022) with the distance metric set to Bray-Curtis. Marginal PERMANOVAs were run the same way, adding the by=“margin” argument. In all cases, we tested for equal dispersion with the betadisper function (equal unless stated otherwise) and ran post-hoc Pairwise Adonis tests in the pairwise.adonis package (v0.4.1; Arbizu, 2017).

We calculated the Shannon-Weiner diversity index with the diversity function in package vegan (Oksanen et al., 2022). The Bray-Curtis dissimilarity index between samples was calculated with the divergence function in the package microbiome (v1.22.0; Lahti & Shetty, 2012) as in Lesser et al. (2016).

To compare means between groups, we ran an analysis of variance (ANOVA) followed by a Tukey post-hoc test. In all cases, we tested the assumption of equal variance between groups with Levene’s test in the package car (v3.1-1; Fox & Weisberg, 2019) and validated the assumption of normality with QQ plots. When the assumption of normality was violated, we ran a Kruskal-Wallis test followed by a post-hoc Wilcoxon test (R Core Team, 2022).

To identify the core *Saccharina* ASVs, we selected all field samples collected at salinity 20 psu or greater and ran an indicator species analysis (IndVal) on non-rarefied data with the function multipatt from the package indicspecies (v1.7.12; Caceres & Legendre, 2009) with 999 permutations, comparing substrate types (*Saccharina*, water, rock, and fake kelp). We used a threshold of 0.7 IndVal statistic, which required ASVs to be enriched and highly prevalent on *Saccharina* compared to other sample types present on a majority of *Saccharina* samples. To calculate the relative abundance of the core across temperature and salinity gradients, we ran linear regression models. We selected the best model by backwards Akaike Information Criterion (AIC) and tested the linear regression assumptions with pots.

Taxa summary plots were generated by calculating the 10 taxa with the greatest relative abundance across all *Saccharina* samples in the field at the order and genus level. We ran linear regression models for each taxa identified as part of the top 10 most abundant taxa in the taxa plot to check for trends in relative abundance along the temperature and/or salinity gradients. We corrected for multiple comparisons with a Benjamini-Hochberg correction (R Core Team, 2022). All plots were made with the ggplot2 (Wickham, 2016) and ggh4x (v0.2.3; Brand, 2022) packages and were saved as pdf files with ggpubr (v0.5.0; Kassambara, 2022a). Text modifications for plot labels were made with the package stringi (v1.7.12; Gagolewski, 2022).

### Reanalysis of published data

Raw data from King et al. (2023) were downloaded from ENA (PRJEB50679) and https://doi.org/10.6084/m9.fgshare.19453889.v1, processed with the dada2 pipeline, and filtered as described above and in their original publication. Core *Saccharina* ASVs were identified as those with a prevalence ≥ 0.8, replicating methods in King et al. (2023) as there are no comparison environmental samples available.

## Results

### Factors shaping the bacterial community of Saccharina in the field

We examined how salinity shapes bacterial community diversity and composition in the field in comparison to temperature on *Saccharina* and in the surrounding environment. We surveyed *Saccharina*-associated bacterial communities biweekly at four sites (Figure 1A) with contrasting salinity (Figure 1C) and temperature (Figure 1D) profiles for two consecutive years during the annual low-salinity event that occurs due to the spring freshet of the Fraser River near Vancouver, Canada. Salinity varied over time and across sites. Salinity varied from 31 psu (typical ocean salinity in the Salish Sea) to a minimum of 10 psu (Figure 1C). Variation in salinity is negatively correlated with temperature, and both salinity and temperature are correlated with seasonality, as measured by Julian day (Figure S1), so we do not include Julian day as a variable in the model to avoid overfitting.

Across the 519 samples in the field dataset, bacterial community composition on *Saccharina*, water, rock, and latex kelp analogues (fake kelp) we installed at our sites in 2022 to assess the biofilm community on substrate with similar morphology are all significantly different from each other by PERMANOVA (Figure S4).

Then, we analyzed the influence of salinity and temperature on *Saccharina*, water, and rock separately using marginal PERMANOVAs nesting within extraction kit. Fake kelp samples were not analyzed further due to low sample size that resulted from loss in the field. We employed a marginal PERMANOVA to assess the unique explanatory power of temperature and salinity, which are highly correlated (Figure S1), so overall variation explained overlaps substantially. Salinity and temperature explain 1.3% and 1.4%, respectively, of unique variation in the bacterial community of *Saccharina* for QIAGEN-extracted sample. Temperature, but not salinity, is a significant explanatory factor (2.2%) for Zymo-extracted samples (Table 1; Figure S5A,D). The rock samples show no significant influence of temperature or salinity on the bacterial community (Table 1; Figure S5B,E). For water samples, water temperature and salinity show the same patterns as for *Saccharina* but explain more unique variation in community composition (Table 1; Figure S5C,F). These results show that the *Saccharina* bacterial community has less turnover than the water bacterial community and more than the communities on rocks across the same environmental conditions.

**Table 1.**
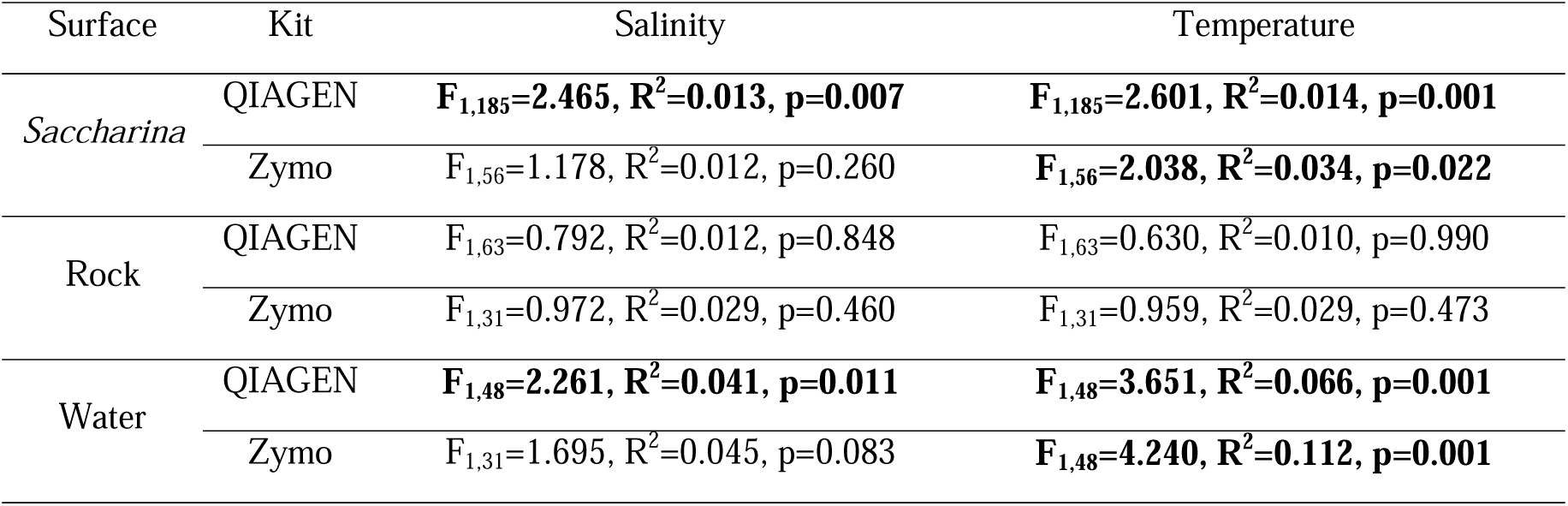
Output of field data marginal PERMANOVA showing the explanatory power marginal explanatory of salinity and water temperature by sample type within the same extraction kit. Statistically significant output is bolded.

| Surface | Kit | Salinity | Temperature |
| --- | --- | --- | --- |
| Saccharina | QIAGEN | **F<sub>1,185</sub>=2.465, R<sup>2</sup>=0.013, p=0.007** | **F<sub>1,185</sub>=2.601, R<sup>2</sup>=0.014, p=0.001** |
|  | Zymo | F<sub>1,56</sub>=1.178, R<sup>2</sup>=0.012, p=0.260 | **F<sub>1,56</sub>=2.038, R<sup>2</sup>=0.034, p=0.022** |
| Rock | QIAGEN | F<sub>1,63</sub>=0.792, R<sup>2</sup>=0.012, p=0.848 | F<sub>1,63</sub>=0.630, R<sup>2</sup>=0.010, p=0.990 |
|  | Zymo | F<sub>1,31</sub>=0.972, R<sup>2</sup>=0.029, p=0.460 | F<sub>1,31</sub>=0.959, R<sup>2</sup>=0.029, p=0.473 |
| Water | QIAGEN | **F<sub>1,48</sub>=2.261, R<sup>2</sup>=0.041, p=0.011** | **F<sub>1,48</sub>=3.651, R<sup>2</sup>=0.066, p=0.001** |
|  | Zymo | F<sub>1,31</sub>=1.695, R<sup>2</sup>=0.045, p=0.083 | **F<sub>1,48</sub>=4.240, R<sup>2</sup>=0.112, p=0.001** |

### Lab salinity treatments differentially induced stress in Saccharina

We tested the influence of salinity on *Saccharina* and associated bacteria using replicated experiments throughout the spring freshet by incubating field-collected *Saccharina* in one of three salinity treatments: 10 psu, 20 psu, and full-strength (31–32 psu; Figure 1E). These are biologically relevant salinity levels for *Saccharina* and are within the range of normal variation experienced at our sites (Figure 1C). We verified that these salinity treatments differentially induced stress in *Saccharina* by measuring four stress-associated kelp phenotypes. We assessed blistering incidence on Day 4 by Fisher’s Exact Test followed by a pairwise Fisher post-hoc test, which shows that our lowest salinity treatment had significantly greater blistering incidence than the other salinity treatments (Figure 2A). The same pattern is observed on Day 1. Experimental round one lasted six days and experimental round two lasted three days. We did not observe any blistering on the last day of these rounds. However, on Day 6 of round one, the *Saccharina* in the 10 psu salinity were green (Figure 2A) and disintegrated when touched, indicating death. This was not observed again throughout the experiment, even when we extended experimental round five to fourteen days, highlighting the importance of adequate biological replication.

We used ANOVA followed by Tukey Post-Hoc tests for the other comparisons. The 10-psu salinity treatment had a significantly lower dry weight to wet weight ratio compared to the other salinity treatments (Figure 2B). The ratio of wet weight on Day 4 compared to Day 0 did not differ by salinity treatment (Figure 2C), and the same results were observed on Day 1. We observed a no significant difference in estimated effective quantum yield on Day 4 (Figure 2D), or Day 1. These results suggest that *Saccharina* is accumulating moisture but not additional biomass in low salinity and remains alive in our lab experiments for the duration of our four-day experiment.

After establishing that our biologically relevant salinity treatments were stressful for *Saccharina*, we assessed the bacterial community composition in the lab. We compared the bacterial community of *Saccharina*, water, and the substrate samples with a PERMANOVA and found that on Day 4, all sample types are significantly different from each other (Figure S4).

Then, we assessed the impact of salinity and experimental round on the bacterial community for each sample type in the lab on Day 4 (Table 2; Figure S5). Our PERMANOVA results follow the same trend as the field data, with the water samples showing the most variation by salinity treatment (Table 2; Figure S5I), followed by *Saccharina* (Table 2; Figure S5G), with the substrate samples (Table 2; Figure S5H) showing no significant change in bacterial community composition by salinity. In our *Saccharina* and water samples, we also see a strong effect of experimental round, showing that the overall effect of salinity in the lab is robust to different starting bacterial communities.

**Table 2.** Output of lab Day 4 PERMANOVA showing the explanatory power salinity and experimental round by sample type. We include the Tukey post-hoc comparisons for salinity treatments when the PERMANOVA showed significant differences between treatments. All three columns of the output for *Saccharina* and water are for the crossed PERMANOVA, while the substrate output is for a PERMANOVA with only salinity. We could not run the crossed PERMANOVA design for the substrate samples because the model is over-fitted due to the smaller sample size of substrate samples (n=14) compared to *Saccharina* (n=26) and water samples (n=27). Statistically significant output is bolded.

### The Saccharina bacterial community is largely stable with a consistent core community

We assessed the stability of the *Saccharina* bacterial community by plotting the most abundant taxa (order and genera) over time and determining whether there is a stable core community on *Saccharina* at the ASV level.

Plotting the most abundant taxa observed on *Saccharina* through the freshet highlights the stability of the *Saccharina* microbiome over time, as much of the community is consistently present (Figure S6A,B). To quantitatively assess the stability of the 10 most abundant taxa, we ran linear regression models followed by Benjamini-Hochberg correction to identify taxa that significantly change in relative abundance over the salinity and temperature gradients (Table S3). At the order level (Figure S6A), there are no significant changes in relative abundance across the salinity or temperature gradient for either extraction kit. At the genus level (Figure S6B), the QIAGEN-extracted samples detect a significant decrease of *Litorimonas* (Caulobacterales) with higher temperatures, whereas the Zymo-extracted samples show a significant increase in *Pseudoalteromonas* (Alteromonadales) relative abundance in lower salinity and higher temperature (Table S3).

We repeated these analyses in the lab with an ANOVA. At the order level (Figure S6C), there is a significant decrease in Caulobacterales in the 10 and 20 psu treatments compared to full-strength salinity (Table S4). A similar pattern is observed at the genus level (Figure S6D), where there is a significant decrease of *Litorimonas* and *Robiginitomaculum* (both Caulobacterales) in the lower salinity treatments (Table S4). These findings, paired with the field data, suggest that Caulobacterales decrease in relative abundance under abiotic stress conditions. We also observed a trend toward increasing Alteromondales at the order level, and *Pseudoaltermonas* (Alteromonadales) at the genus level in low salinity, which are opportunistic colonizers.

We identified the core bacterial community on *Saccharina* when the salinity was relatively high (20 psu or greater). We excluded lower salinity samples when defining the core because these conditions are likely to be stressful to the *Saccharina* populations studied here based on the results of the lab experiments (Figure 2). We used indicator species analysis with a threshold of 0.7 IndVal value to identify ASVs that were enriched on *Saccharina* compared to water and rock and at high frequency on *Saccharina* when ocean salinity was greater or equal to 20 psu. We identified six core ASVs (Table S5).

Then, we investigated the distribution of core bacteria ASV on *Saccharina* across much larger spatial scales by comparing our data to a study performed in the United Kingdom (UK) from September and August 2015 (King et al., 2023). The UK study did not include environmental comparison samples, so core bacteria were defined solely based on a prevalence of 0.8 or higher; this resulted in 27 core ASVs. Comparing the core ASVs between the two datasets, 4/6 core ASVs detected in our BC population are also found to be in association with *Saccharina* in the UK, with ASV7 (*Robiginitomaculum* sp.) being a core ASV in both datasets (Table S5). Nineteen of the twenty-seven core ASVs in the UK dataset are present in our *Saccharina* samples, including ASV7. However, many of these ASVs are present at low prevalence, highlighting the importance of including environmental samples in these types of analyses. When we compare the BC and UK datasets at the genus level, we find that all genera that include core *Saccharina* ASVs are present in the other dataset, indicating that functional redundancy within a genus, paired with other factors, is likely responsible for differences in ASV-level overlap (Table S5). Overall, these comparisons show that, to a large extent, the *Saccharina* bacterial community is consistent across global scales.

### Is bacterial community change due to turnover or destabilization of the community?

We asked whether changes in the bacterial community associated with *Saccharina* are consistent with a shift to a distinct low-salinity community (turnover) or destabilization of the community. Destabilization in stressful conditions is predicted to occur for hosts that have an otherwise stable microbiome, with the assumption that stress reduces host filtering. *Saccharina* has a stable bacterial community, as discussed above (Figure S6, Table S5). The Anna Karenina Principle (AKP) specifically predicts increased community dissimilarity (beta diversity) in stressful conditions. Additionally, we predicted that increased beta diversity would be accompanied by increased alpha diversity and reduced relative abundance of core taxa if host filtering is diminished in low-salinity stress, or high-temperature stress in the field. The lab experiment allows us to isolate the influence of salinity. Moreover, two years of field-survey data allow us to determine whether the patterns observed in the lab are consistent with observed dynamics on natural populations. Interpretation of patterns in the field is complicated by the fact that decreasing salinity in the field is correlated with seasonality patterns, generally, and increasing temperature in particular (Figure S1), which is likely an additive stressor for kelp in the Salish Sea.

We tested for turnover in the bacterial community of *Saccharina* by asking if there is a distinct core community in low-salinity conditions in the field or the lab. We identified the core bacterial community of *Saccharina* from low-salinity field samples (below 20) using the same methods as above. We found that the core *Saccharina* bacterial community from low-salinity field samples is comprised of 10 ASVs (Table S5). This includes five of the six ASVs that are part of the high-salinity field core; the sixth high-salinity core ASV (ASV15) is also present in low salinity, with an IndVal stat of 0.62, slightly below the 0.7 IndVal threshold, though still common on *Saccharina* in low-salinity conditions (Table S5). The five other ASVs that are core in low salinity (salinity <20) but not high salinity (salinity ≥ 20) in the field have IndVal stats ranging from 0.67 to 0.45 in the high-salinity field samples, indicating again, that these ASVs are commonly associated with *Saccharina* in higher salinity conditions (Table S5). These data show that although the ASVs that meet the threshold of IndVal stat 0.7 differ somewhat between the high and low-salinity field samples, all of the core ASVs are present across the salinity gradient and there is not a distinct core bacterial community in low salinity in the field.

The persistence of the *Saccharina* core community across the salinity gradient is also supported lab data showing that the core ASVs are maintained in the lab and at high prevalence across all salinity treatments (Table S5). We observe 100% prevalence for five of the six high salinity core ASVs and 89% prevalence for ASV15 at 20 and 30 psu. All core ASVs persist at the lowest salinity, though four decline in prevalence (50% to 87%; Table S5). Overall, these data paint a picture of a stable microbial community on *Saccharina* characterized by distinct core bacteria that persist in the lab and at low salinity with no evidence of turnover to a distinct and consistent low-salinity community.

Next, we tested our prediction that destabilization would lead to reduced abundance of the core microbiome and increased dissimilarity and alpha diversity in stressful conditions. Low salinity and high temperature represent stressful conditions in the field, while the lab experiment isolated the influence of low-salinity stress. For the field data, we ran linear regression models including temperature and salinity, nested within theextraction kit and selected the best model by backwards stepwise model selection using AIC. For the lab, we used ANOVAs to compare core relative abundance, alpha diversity, and dissimilarity across treatments.

#### Relative abundance of core ASVs

We find that the best model for the relationship between the total relative abundance of the six *Saccharina* core ASVs and temperature and salinity (F_5,241_=30.13, adjusted-R^2^=0.372, p<0.001) includes both salinity and temperature. Within this model, for the QIAGEN-extracted samples both salinity (t=2.528, p=0.0121; Figure 3A) and temperature (t=-2.110, p=0.036; Figure 3B) are significant, but for the Zymo-extracted samples, neither salinity (t=0.466, p=0.642; Figure 3A) nor temperature (t=-1.250, p=0.213; Figure 3B) are significant. We repeated this analysis with thresholds 0.5 to 0.8 IndVal to ensure the patterns observed were a robust choice of threshold for defining the core; they were (Figure S7). In the lab by ANOVA (F_2,24_=16.05, p<0.001) the core decreases in relative abundance in the 10 and 20 psu treatments compared to the full-strength (p≤0.007; Figure 3C). The lab and field data show that core *Saccharina* taxa are maintained at relatively high prevalence at all salinities in the field and in the controlled lab environment. These data support the broad patterns of stability in the *Saccharina* bacterial community observed in the taxa plots (Figure S6). The decreased relative abundance of the core observed on *Saccharina* both in the lab and field are consistent with our predictions for community destabilization in response to abiotic stress—temperature (field only) and salinity (lab and field).

**Figure 3.**
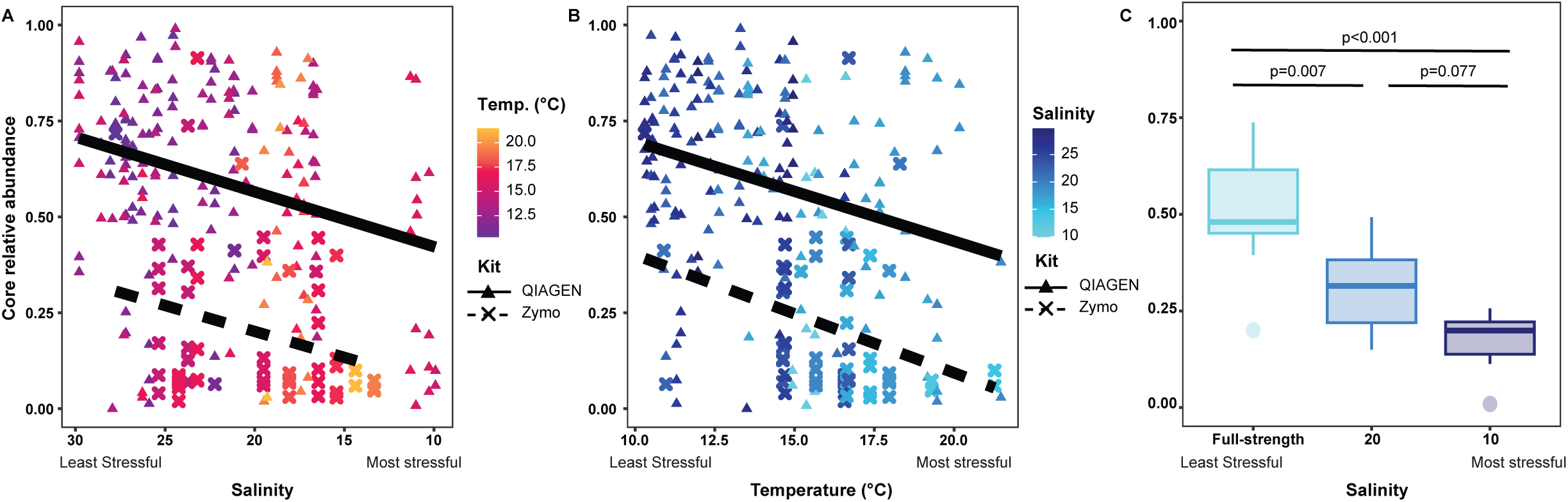
Relative abundance of core ASVs in the field (A, B) and in the lab on Day 4 (C) on *Saccharina*. Plots A and B show the same samples, but arranged along the salinity (A) and temperature (B) gradients observed in the field across both years. The colour gradients in panels A and B show the abiotic gradient not plotted on the x-axis. Point and line shape show the extraction kit used to extract the samples. In C, the Tukey posy-hoc comparison p-values for the ANOVA are shown. Note, all three x-axes are arranged from least to most stressful abiotic condition.

#### Alpha diversity

The best model for alpha diversity of the field *Saccharina* samples includes temperature but not salinity (Figure 4A, Table S6); there is a significant increase in alpha diversity for the Zymo-extracted samples from warmer water, but no change in the QIAGEN-extracted samples. Since the Zymo-extracted samples include few time points, this trend may not be robust. The best model for the rock samples includes only the extraction kit, showing no pattern in alpha diversity by salinity or temperature (Figure 4B, Table S6). Lastly, the best model for water samples includes salinity but not temperature, with the QIAGEN-extracted samples showing increased diversity in lower salinity (Figure 4C, Table S6). We then assessed the effect of salinity on alpha diversity in the lab on Day 4 by ANOVA and found that for all of the sample types (*Saccharina*, substrate, water) there are no significant differences in alpha diversity by salinity treatment (Figure 4D,E,F, Table S6). Overall, there is scant evidence for a relationship between alpha diversity in the *Saccharina* microbiome and stressful conditions.

**Figure 4.**
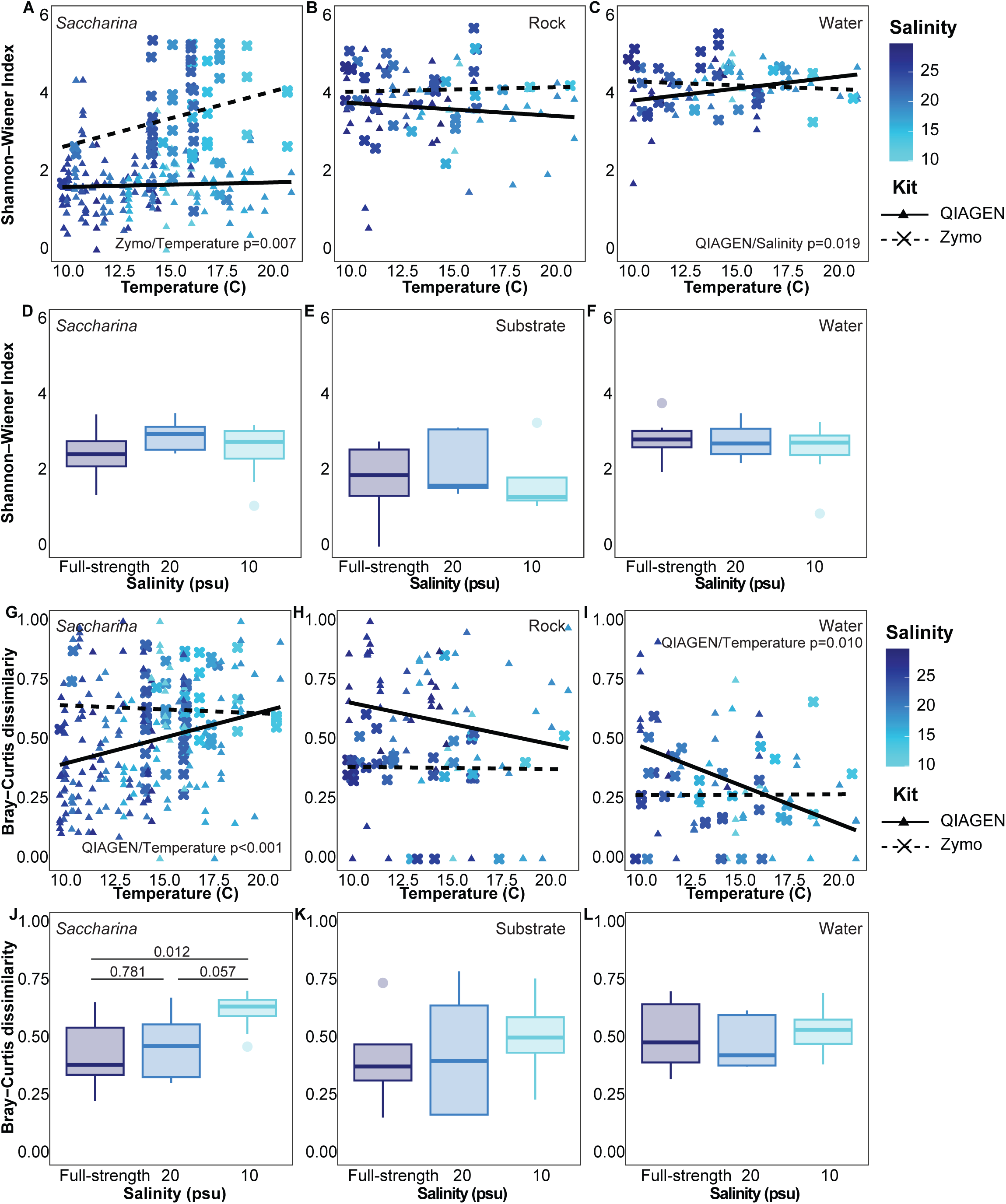
The Shannon-Weiner diversity index for the field (A,B,C) and lab Day 4 (D,E,F) samples along with the Bray-Curtis dissimilarity index of the field (G,H,I) and lab Day 4 (J,K,L) samples. Beta diversity is calculated within the same substrate type either by comparing samples within the same sampling site visit (field) or a salinity treatment (lab). The same sample types are arranged in columns and indicated in the panels. For the field samples, the point colour represents salinity and the shape (point and line) indicates the extraction kit. For all panels, the full statistical output is in Table S6, but we indicate the significant factors in the nested linear regression model (field) or Tukey post-hoc test if the ANOVA showed significant differences between groups (lab). Note, all axes are ordered from least stressful to most stressful, with the field sampling plotted along the temperature gradient while the lab samples are plotted by salinity treatment.

#### Community dissimilarity

We repeated the same process to assess changes in dissimilarity. The best model for field *Saccharina* includes temperature only, with the QIAGEN-extracted sample showing a significant increase in dissimilarity in higher temperatures (Figure 4G, Table S6). The best model for the rock samples includes the extraction kit only, again showing no pattern by abiotic conditions (Figure 4H, Table S6). The best model for the water samples includes temperature and salinity, with QIAGEN-extracted showing a significant decrease in beta diversity in higher temperatures (Figure 4I, Table S6). In the lab, the dissimilarity of the *Saccharina* bacterial communities in the 10 psu treatment is significantly higher compared to the 20 psu and full-strength salinities (Figure 4J, Table S6), but there is no significant change in the water and substrate communities (Figure 4K,L, Table S6). Overall, the results in the field and lab show that *Saccharina*, but not water or rock, show the predicted AKP pattern of increased dissimilarity in stressful conditions. In the field, temperature has a stronger effect than salinity on community dissimilarity. Temperature and salinity are likely additive stressors in nature. In summary, our data support the predicted AKP pattern of increased beta diversity associated with abiotic stress and we find a consistent decrease in core relative abundance but no relationship with alpha diversity in stressful conditions.

## Discussion

The predicted (Bindoff et al., 2019; Filbee-Dexter et al., 2019) and ongoing die back of kelp forests (Bindoff et al., 2019; Christie et al., 2019; Davis et al., 2022) due to climate change represents a global concern because of the environmental, economic, and cultural value that these foundation species provide (Bindoff et al., 2019; Eger et al., 2023). We find that salinity influences the bacterial community of *Saccharina,* in agreement with other macroalgal studies in the lab (*Agarophyton*; Saha et al. (2020) and *Fucus*; Stratil et al. (2014)) and in the field (*Ulva*; van der Loos et al. (2023) and *Nereocystis*; Weigel & Pfister (2019)) showing that salinity influences the bacterial community associated with macroalgae.

A change in the bacterial community composition can influence host health in a variety of different ways. In some cases, a short-term change in the bacterial community allows the host to better tolerate stressful abiotic conditions (high temperatures, Baldassarre et al., 2022; Ziegler et al., 2017). Heat stress in the kelp *Ecklonia* results in bacterial community turnover, but no interaction between microbial community and the host stress response (Vadillo Gonzalez et al., 2024). This represents a directional turnover in the bacterial community, rather than a destabilized community. In other cases, host-associated bacteria are variable across non-stressful conditions including localities (Seagrass, Adamczyk et al., 2022) and natural salinity gradients (*Ulva*, van der Loos et al., 2022). In other cases, a destabilization of the bacterial community— characterized by increased beta diversity and declining abundance of characteristic taxa—is associated with host stress. A destabilized bacterial community associated with host stress (usually high temperature) has been observed in corals (McDevitt-Irwin et al., 2017) and on some sponges (Pita et al., 2018).

Our lab experiments at biologically relevant salinity levels show that low salinity induces stress in *Saccharina*, which allows us to investigate the bacterial community changes associated with stress, and to determine whether the community changes observed in low salinity are the results of turnover or destabilization. In the case of turnover (option 1), we expect to see a new, stable, bacterial community in low salinity. In the case of destabilization (option 2), we expect increased community dissimilarity (as predicted by AKP, Zaneveld et al., 2017) along with a reduction of core bacteria and increased alpha diversity, which indicated decreased host filtering in low salinity.

Our lab and field data both support option 2: *Saccharina*-associated bacterial community is destabilized in low-salinity conditions. In both the lab and the field, the relative abundance of core ASVs decreases in lower salinity (Figure 3), but these core ASVs are not replaced by a distinct and consistent low-salinity community (Table S5). At the genus and order level, we find that most taxa are present at similar relative abundances at all salinities, also showing no clear evidence of turnover (Figure S6, Table S3, Table S4). Additionally, our data show increased dissimilarity in low salinity (Figure 4), supporting the AKP prediction (Zaneveld et al., 2017), likely due to a decrease in host filtering. We find no associated trends in alpha diversity with salinity (Figure 4), showing that increased dissimilarity is not explained by trends in alpha diversity. Our study was not designed to test if the changes we observed in the bacterial community of *Saccharina* are beneficial or not to kelp’s ability to tolerate low salinity. Testing this would require controlling for host genetics (as performed with anemones by Baldassarre et al. (2022) and with corals by Ziegler et al. (2017) and directly manipulating the bacterial community (as by Dittami et al. (2016) with *Ectocarpus* and Santoro et al. (2021) with coral).

Our study highlights the importance of pairing field observations, which provide a more holistic view of the system, with lab experiments, which permit the manipulation of a single variable (salinity) without confounding effects of other covariates present in the field (as suggested by Trevathan-Tackett et al., 2020). Here, salinity is strongly correlated with water temperature during the spring freshet in the Fraser River Estuary and temperature appears to be a stronger factor shaping the bacterial community of *Saccharina* according to our field data. Temperature explains slightly more variation in bacterial community composition (Table 1). Temperature is a significant predictor of community dissimilarity in the multivariate model assessing the relationship between dissimilarity and abiotic stress (Figure 4), but salinity is not. Thus, the lab experiment conducted at local biologically relevant salinity levels (Figure 1) allows us to conclude that salinity can produce the changes in the *Saccharina* bacterial community observed in the field.

## Abbreviations

UBC: University of British Columbia
IndVal: indicator species analysis
ENA: European Nucleotide Database
PERMANOVA: Permutational multivariate analysis of variance
AKP: Anna Karenina Principle
AIC: Akaike Information Criterion
BC: British Columbia
UK: United Kingdom

## Data Availability

Raw reads for this study are available on the European Nucleotide Database (ENA) under project PRJEB60884. All code and metadata are available on Borealis https://borealisdata.ca/dataset.xhtml?persistentId=doi:10.5683/SP3/ILQ9UJ.

## Contributions

Siobhan Schenk: conceptualization (equal), data collection (equal), writing (equal), editing (equal), data analysis (equal).

Connor G. Wardrop: data collection (equal), editing (equal).

Laura W. Parfrey: conceptualization (equal), data collection (equal), writing (equal), editing (equal), data analysis (equal).

## Acknowledgements

We thank E. Adamczyk, G. Ainsworth-Cruickshank, V. Billy, O. Moss, and V. Supratya for their help with fieldwork, B. Segovia and G. Lajoie for their help with data analysis, R. d’Entremont for editing the latest version of the manuscript, the UBC staff who helped us with the lab experiment, in particular J. Trat and J. Ng, and Hakai for their expertise with Illumina Next Generation Sequencing.

## Supplemental Text Figure and Table Legends

**Figure S1.**
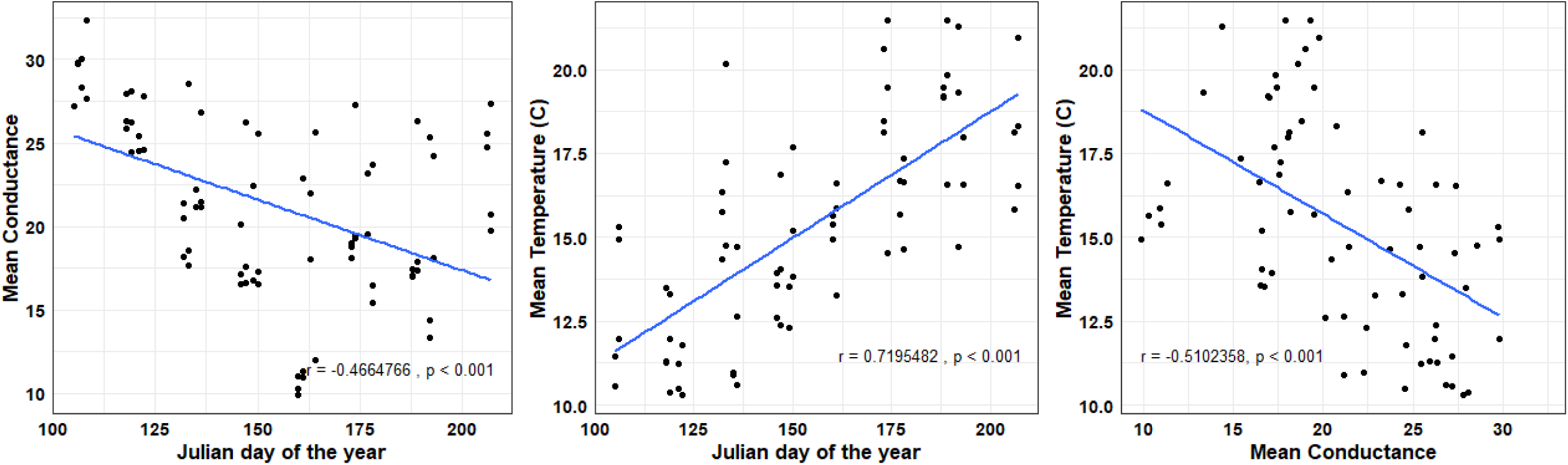
Dot plots with regression line showing the correlation between conductance (µS/cm; referred to as salinity in the text), temperature (°C), and Julian day for 2021 and 2022. Results of Pearson’s Correlation Test indicated in the plot area.

**Figure S2.**
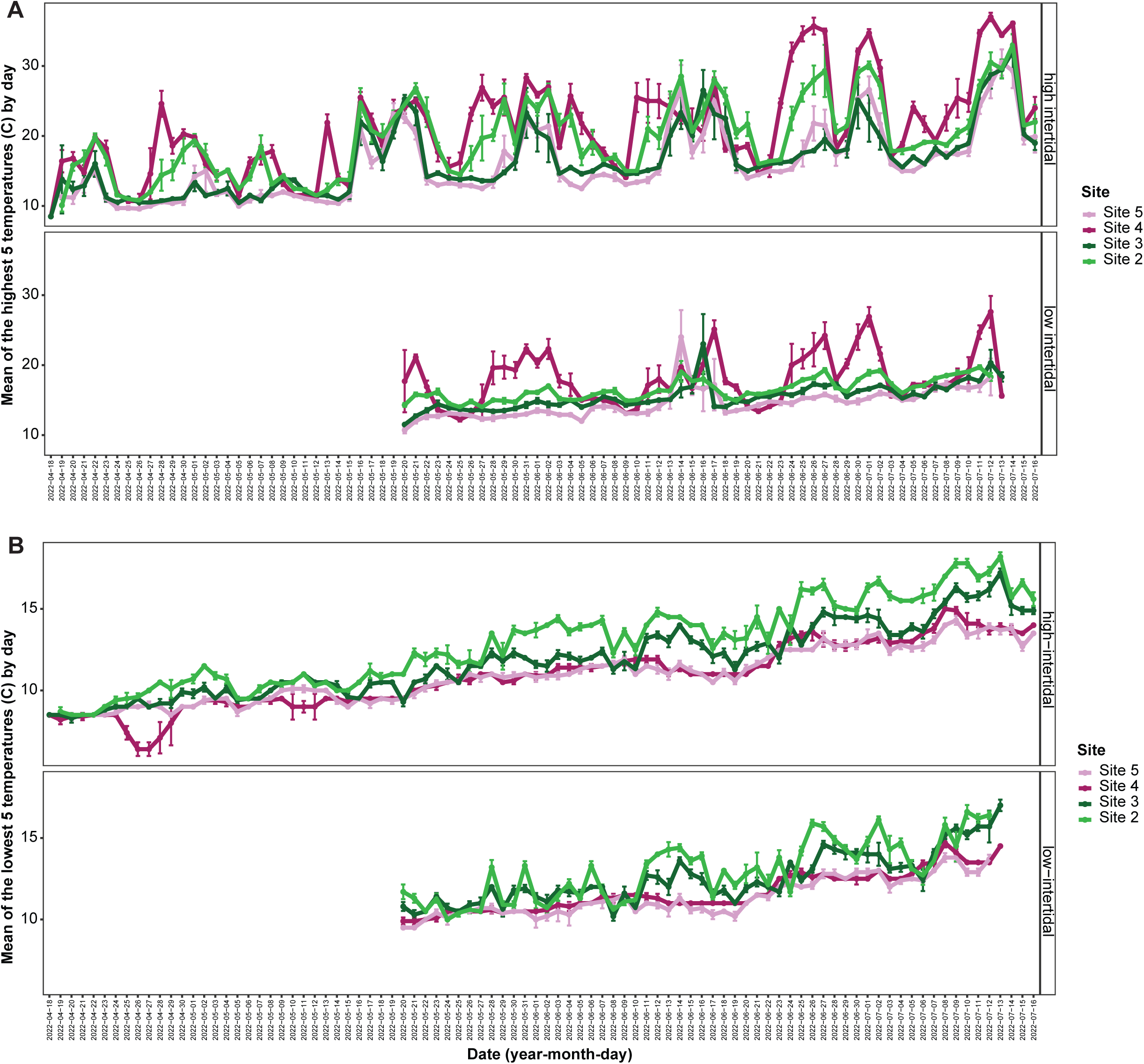
The mean 5 (A) highest or (B) lowest daily recorded temperatures (°C) by each temperature logger. Error bars are +/-1 standard deviation.

**Figure S3.**
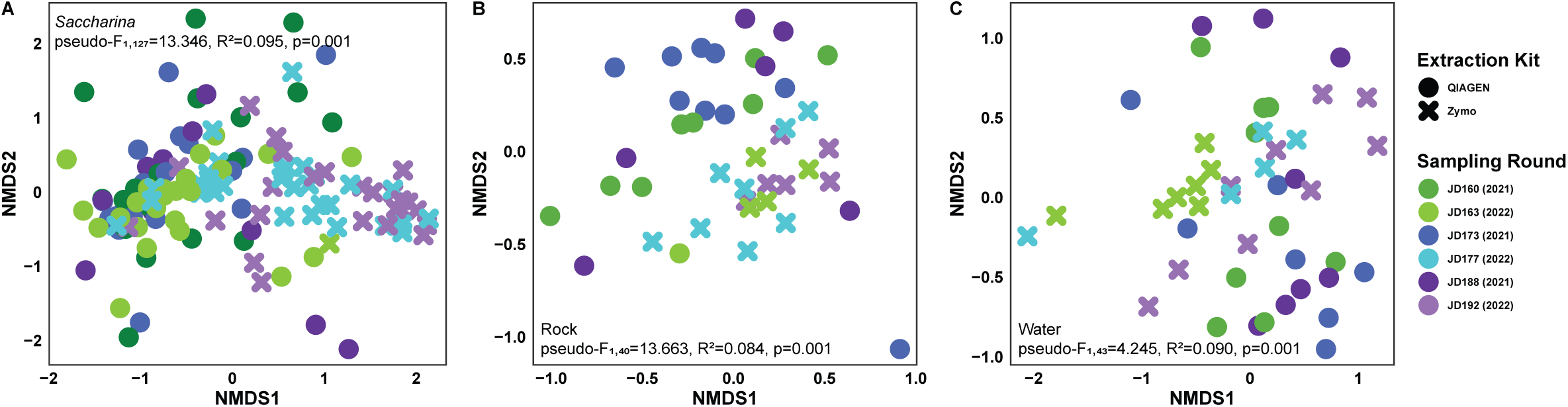
NMDS plots showing field A) *Saccharina*, B) rock, and C) water samples. Point colour represents the Julian day sampling round (Figure 1B), with equivalent rounds being the same colour. Darker shades are 2021 samples and lighter colours are samples from 2022. Point shape represents the extraction kit used to extract that sample. PERMANOVA results comparing extraction kits for each sample type are listed in the corresponding panel.

**Figure S4.**
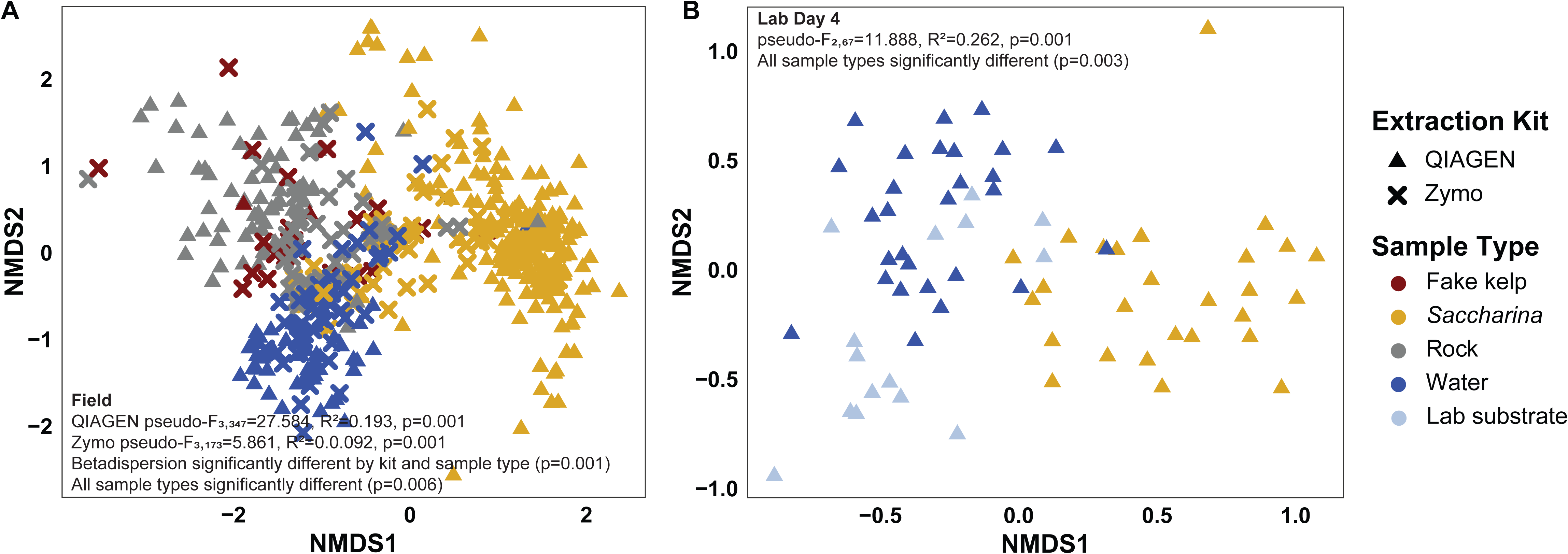
NMDS plots showing each sample type (colour) and the extraction kit used (shape) for A) both years of field samples and B) lab Day 4 samples. PERMANOVA outputs are in the corresponding panels.

**Figure S5.**
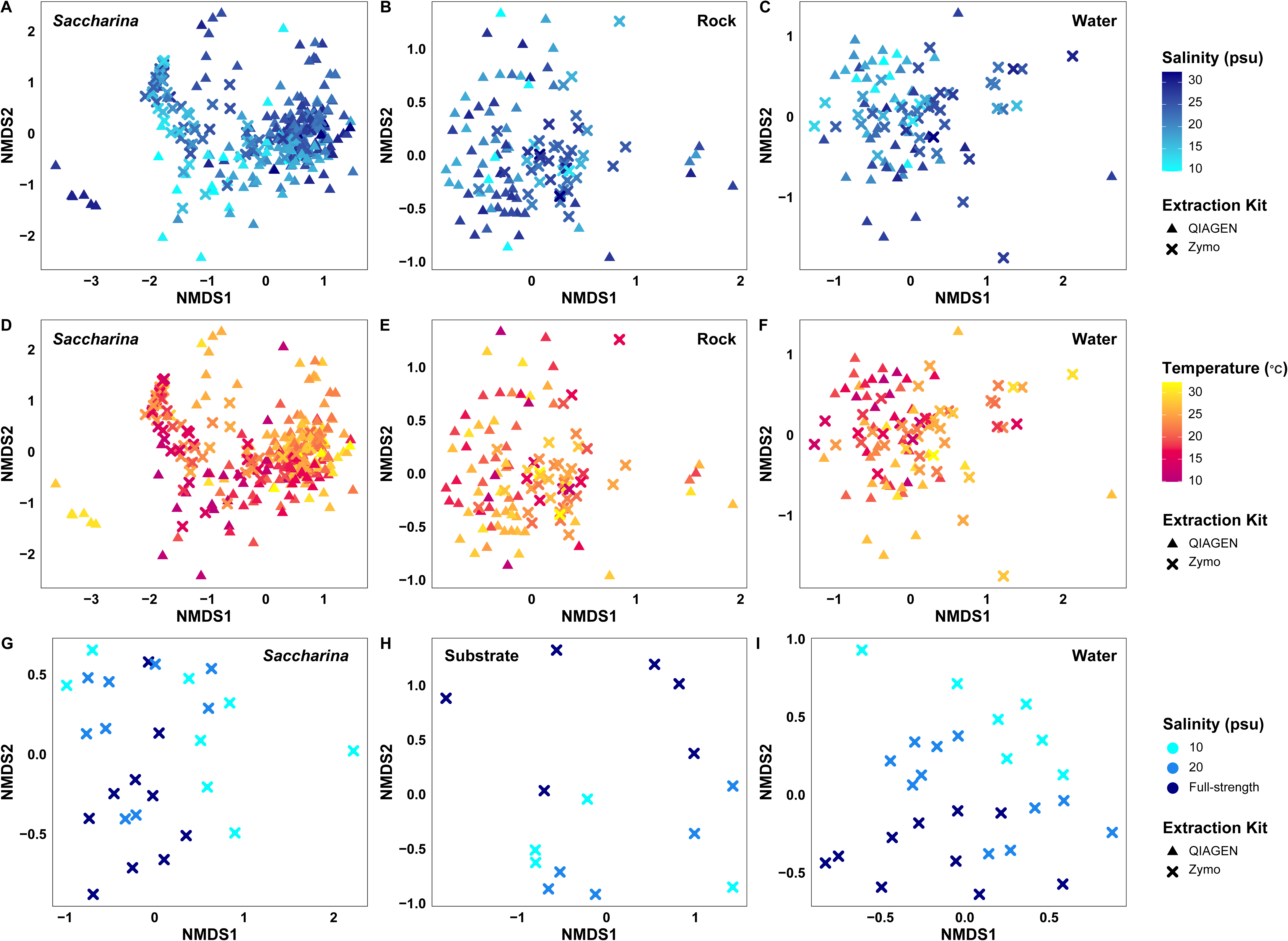
NMDS plots by sample type showing the distribution of samples across the salinity (A:C,G:H) and temperature (D:F) gradients in the study. Panels A to F show field samples, while panels G to I show lab Day 4 samples. Sample types are in the same column, while the same abiotic gradient is shown across rows. Point shapes indicate the extraction kit used.

**Figure S6.**
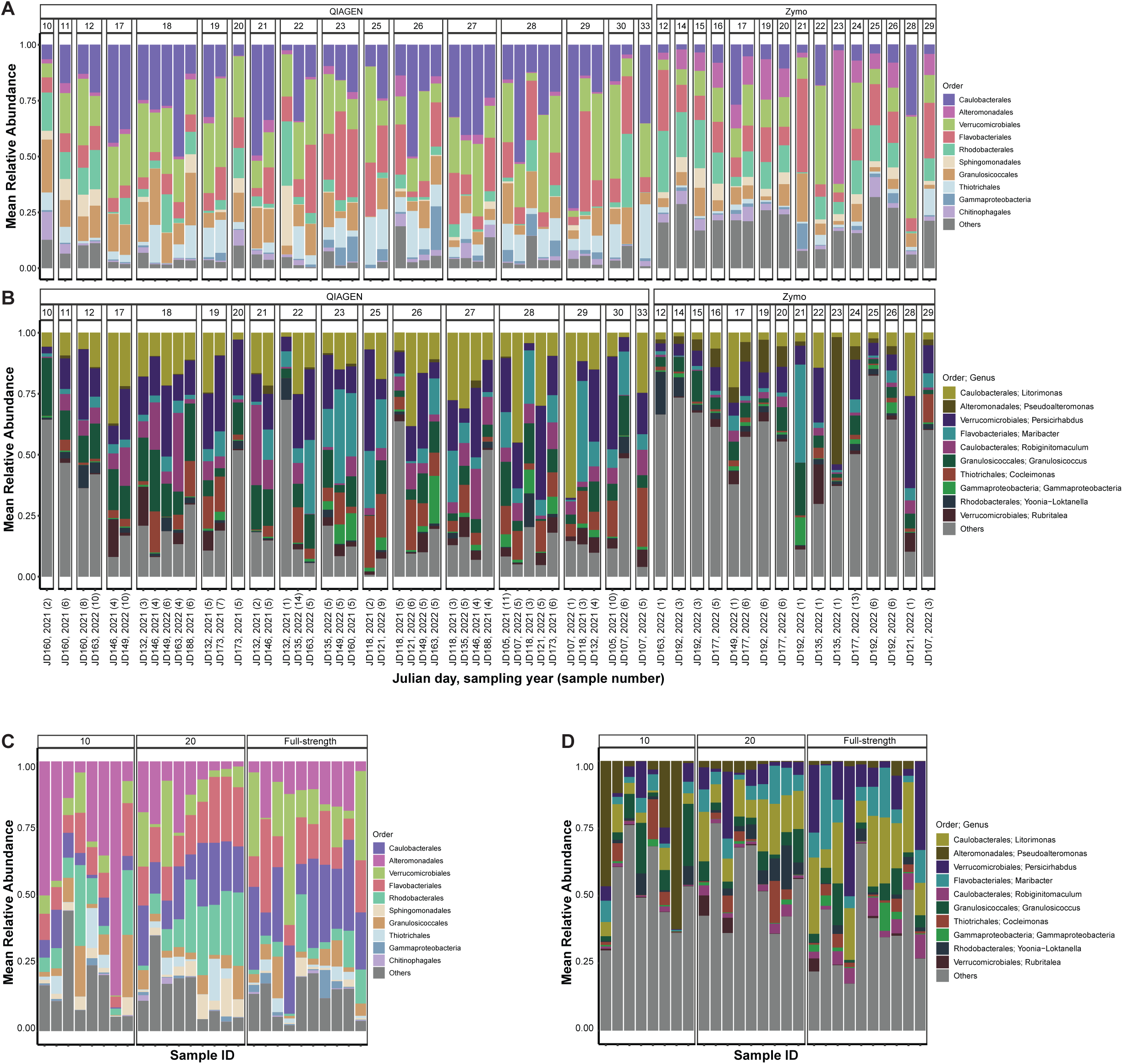
The relative abundance of the 10 most abundant orders (A,C) or genera (B,D) in the field samples. Field samples (A,B) are separated by extraction kit and salinity on the sampling day (facets) and the per-sample relative abundance is averaged across all samples from the same sampling day. Sample numbers are indicated in parentheses. (C,D) Each lab sample is shown (not averaged) and show the 10 most abundant taxa as identified in the field data. Colours are consistent between the field and lab Day 4 plots. Tables include the output of models comparing the relative abundance of taxa in the field (Table S3) and lab Day 4 (Table S4).

**Figure S7.**
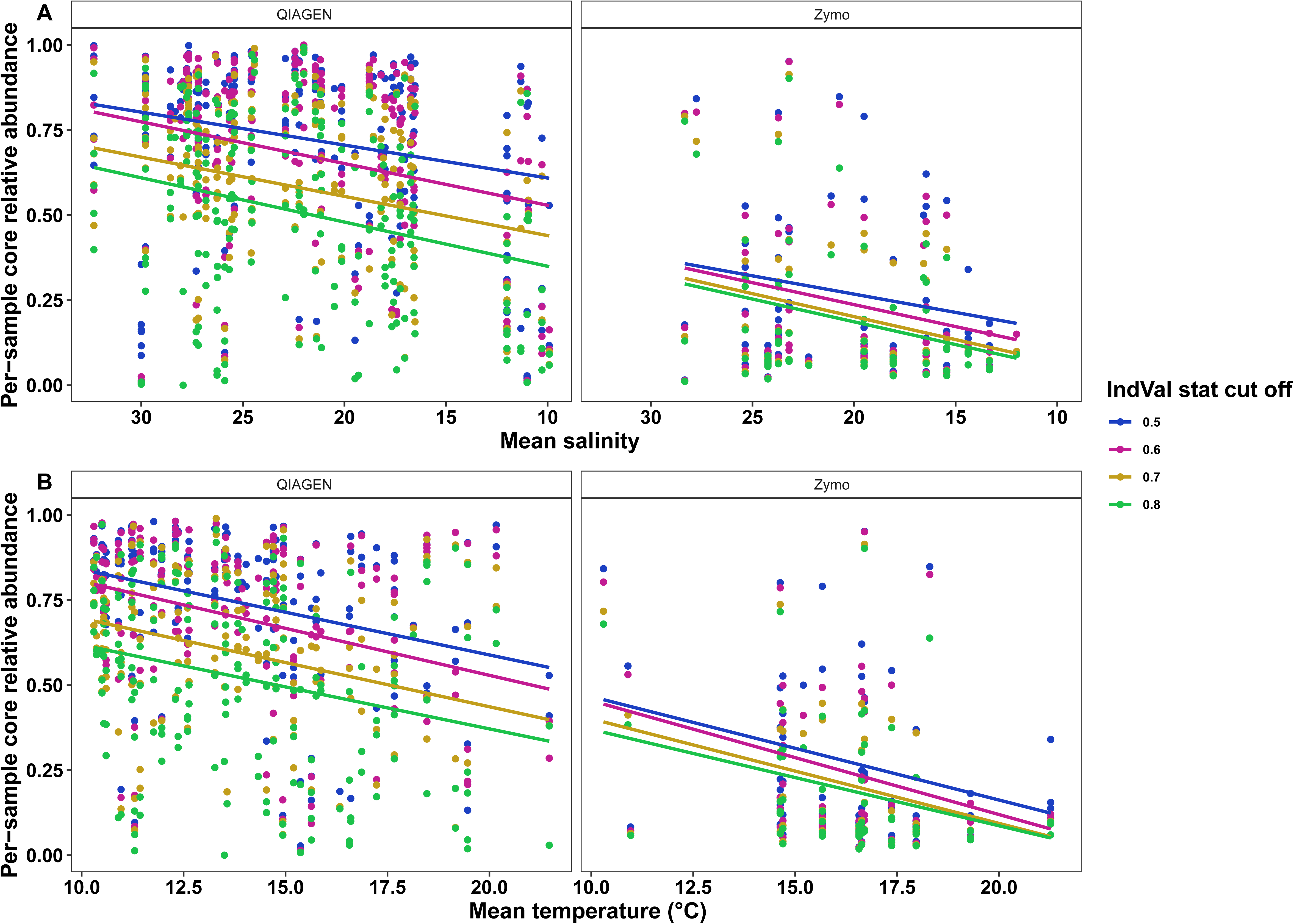
Scatter plot of the per-sample relative abundance of core ASVs on *Saccharina* in the field along the A) salinity or B) temperature gradient in the field arranged from the least stressful on the left to the most stressful on the right. Facets separate the samples by extraction kit and colours represent the different IndVal stat threshold values, meaning each sample is plotted for each IndVal stat value.

**Table S1.** Shipping dates and associated salinity of the full-strength seawater for each lab experimental round.

| Experimental round number (start date) |  |  | New water shipment date | Full-strength salinity |
| --- | --- | --- | --- | --- |
| 1 (April 19, 2022) |  |  | April 13, 2022 | 32 |
| 2 (May 3, 2022) |  |  | April 13, 2022 | 32 |
| 3 (May 17, 2022) |  |  | April 13, 2022 | 32 |
| 4 (May 31, 2022) |  |  | April 13, 2022 | 32 |
| 5 (June 14, 2022) |  |  | June 10, 2022 | 31 |
| Surface | Kit | Salinity | Experimental Round | Salinity*Experimental Round |
| Saccharina | Zymo | **F<sub>2,9</sub>=3.363, R<sup>2</sup>=0.166, p=0.002** | **F<sub>5,9</sub>=2.692, R<sup>2</sup>=0.331, p=0.001** | F<sub>10,9</sub>=1.145, R<sup>2</sup>=0.281, p=0.266 |
|  |  | 10 vs 20 p=0.096 |  |  |
|  |  | **10 vs full-strength p=0.003** |  |  |
|  |  | 20 vs full-strength p=0.294 |  |  |
| Substrate | Zymo | F<sub>2,12</sub>=1.188, R<sup>2</sup>=0.165, p=0.264 | Not run | Not run |
| Water | Zymo | **F<sub>2,11</sub>=4.626, R<sup>2</sup>=0.186, p=0.001** | **F<sub>6,11</sub>=3.273, R<sup>2</sup>=0.394, p=0.001** | F<sub>8,11</sub>=1.241, R<sup>2</sup>=0.199, p=0.113 |
|  |  | 10 vs 20 p=0.189 |  |  |
|  |  | **10 vs full-strength p=0.003** |  |  |
|  |  | **20 vs full-strength p=0.039** |  |  |
| 6 (June 28, 2022) |  |  | June 10, 2022 | 31 |
| 7 (July 12, 2022) |  |  | June 10, 2022 | 31 |
| 8 (July 27, 2022) |  |  | July 19, 2022 | 32 |

**Table S2.** Number of field bacterial samples that were sequenced per sample source, year, and sample type. Also, the sample number of the lab 8 experimental rounds, showing the number of samples per lab experimental day. Note, all experimental rounds included three salinity levels (10 psu, 20 psu, and full-strength), with two replicate aquaria per salinity level. There were three *Saccharina* per aquaria in all instances except experimental round 1, where one of the 10 psu salinity aquaria only had two *Saccharina*.

**Table S3.** Output of linear regression model comparing the per-sample relative abundance of each of the 10 most abundant orders and genera in the field in field *Saccharina* samples. Models were run individually for each taxa and were nested within the extraction kit, where trends in taxa relative abundance by salinity and temperature were assessed. P-values were adjusted for multiple comparisons with a Benjamini-Hochberg correction.

**Table S4.** Output of ANOVA model comparing the per-sample relative abundance of each of the 10 most abundant orders and genera in the field in the Day 4 lab *Saccharina* samples. ANOVAs were run individually for each taxa and p-values were adjusted for multiple comparisons with a Benjamini-Hochberg correction.

**Table S5.** Table showing the output of IndVal analysis for the field, lab, and King datasets. Taxonomy information for each ASV is provided in columns A to I. The sample type to which the ASV was assigned (*Saccharina*, water, substrate), the IndVal statistic, the specificity, and prevalence of the ASV for the assigned substrate are indicated in columns J to M (salinity ≥20) and N to P (salinity <20). Columns R to U indicate the prevalence of the ASV in the lab by salinity (R to T) and in the King data (U). Column V indicates if the ASV was assigned as core in high (salinity ≥20) and/or low salinity (salinity <20). The UK core can be identified in column U (Prevalence_King), with a prevalence of ≥ 0.8.

**Table S6.** Output of the best nested linear regression model by backwards AIC for the field or ANOVA for the lab Day 4 samples. From left to right, the columns indicate if the samples analyzed are from the field or lab, if the model is analyzing alpha (Shannon-Weiner) or beta diversity (Bray-Curtis Dissimilarity), the type of mode, the sample type analyzed, the output of the overall model, the output for each factor in the model (field data only), and the output of the Tukey post-hoc test (lab only).

## References

Adamczyk, E. M., O’Connor, M. I., & Parfrey, L. W. (2022). Seagrass (Zostera marina) transplant experiment reveals core microbiota and resistance to environmental change. Molecular Ecology, 31(19), 5107–5123. 10.1111/mec.16641

Arbizu, P. M. (2017). pairwiseAdonis: Pairwise Multilevel Comparison using Adonis.

Baldassarre, L., Ying, H., Reitzel, A. M., Franzenburg, S., & Fraune, S. (2022). Microbiota mediated plasticity promotes thermal adaptation in the sea anemone Nematostella vectensis. Nature Communications, 13(1), 3804. 10.1038/s41467-022-31350-z

Bindoff, N. L., Cheung, W. W. L., Kairo, J. G., Arístegui, J., Guinder, V. A., Hallberg, R., Hilmi, N., Jiao, N., O’Donoghue, S., Suga, T., Acar, S., Alava, J. J., Allison, E., Arbic, B., Bambridge, T., Boyd, P. W., Bruggeman, J., Butenschön, M., Chávez, F. P., … Whalen, C. (2019). Changing Ocean, Marine Ecosystems, and Dependent Communities. Marine Ecosystems.

Bollen, M., Pilditch, C. A., Battershill, C. N., & Bischof, K. (2016). Salinity and temperature tolerance of the invasive alga Undaria pinnatifida and native New Zealand kelp: Implications for competition. Marine Biology, 163(9), 194. 10.1007/s00227-016-2954-3

Brand, T. van den. (2022). ggh4x: Hacks for “ggplot2.” https://CRAN.R-project.org/package=ggh4x

Caceres, M. D., & Legendre, P. (2009). Associations between species and groups of sites: Indices and statistical inference. In Ecology. http://sites.google.com/site/miqueldecaceres/

Callahan, B. J., McMurdie, P. J., Rosen, M. J., Han, A. W., Johnson, A. J. A., & Holmes, S. P. (2016). DADA2: High-resolution sample inference from Illumina amplicon data. Nature Methods, 13(7), Article 7. 10.1038/nmeth.3869

Chen, M. Y., & Parfrey, L. W. (2018). Incubation with macroalgae induces large shifts in water column microbiota, but minor changes to the epibiota of co occurring macroalgae. Molecular Ecology, 27(8), 1966–1979. 10.1111/mec.14548

Christie, H., Andersen, G. S., Bekkby, T., Fagerli, C. W., Gitmark, J. K., Gundersen, H., & Rinde, E. (2019). Shifts Between Sugar Kelp and Turf Algae in Norway: Regime Shifts or Fluctuations Between Different Opportunistic Seaweed Species? Frontiers in Marine Science, 6, 72. 10.3389/fmars.2019.00072

Davis, K. (2022). Factors structuring microbial communities on marine foundation species [University of British Columbia]. 10.14288/1.0413811

Davis, T. R., Larkin, M. F., Forbes, A., Veenhof, R. J., Scott, A., & Coleman, M. A. (2022). Extreme flooding and reduced salinity causes mass mortality of nearshore kelp forests. Estuarine, Coastal and Shelf Science, 275, 107960. 10.1016/j.ecss.2022.107960

Davison, I. R., & Pearson, G. A. (1996). Stress Tolerance in Intertidal Seaweeds. Journal of Phycology, 32(2), 197–211. 10.1111/j.0022-3646.1996.00197.x

Dittami, S. M., Duboscq-Bidot, L., Perennou, M., Gobet, A., Corre, E., Boyen, C., & Tonon, T. (2016). Host—microbe interactions as a driver of acclimation to salinity gradients in brown algal cultures. The ISME Journal, 10(1), 51–63. 10.1038/ismej.2015.104

Egan, S., Harder, T., Burke, C., Steinberg, P., Kjelleberg, S., & Thomas, T. (2013). The seaweed holobiont: Understanding seaweed—bacteria interactions. FEMS Microbiology Reviews, 37(3), 462–476. 10.1111/1574-6976.12011

Eger, A. M., Marzinelli, E. M., Beas-Luna, R., Blain, C. O., Blamey, L. K., Byrnes, J. E. K., Carnell, P. E., Choi, C. G., Hessing-Lewis, M., Kim, K. Y., Kumagai, N. H., Lorda, J., Moore, P., Nakamura, Y., Pérez-Matus, A., Pontier, O., Smale, D., Steinberg, P. D., & Vergés, A. (2023). The value of ecosystem services in global marine kelp forests. Nature Communications, 14(1), Article 1. 10.1038/s41467-023-37385-0

Eppley, R. W., & Bovell, C. R. (1958). Sulfuric Acid in Desmarestia. Biological Bulletin, 115(1), 101–106. 10.2307/1539096

Filbee-Dexter, K., Wernberg, T., Fredriksen, S., Norderhaug, K. M., & Pedersen, M. F. (2019). Arctic kelp forests: Diversity, resilience and future. Global and Planetary Change, 172, 1–14. 10.1016/j.gloplacha.2018.09.005

Fox, J., & Weisberg, S. (2019). An R Companion to Applied Regression (Third). Sage. https://socialsciences.mcmaster.ca/jfox/Books/Companion/

Gagolewski, M. (2022). stringi: Fast and portable character string processing in R. Journal of Statistical Software, 103(2), 1–59. 10.18637/jss.v103.i02

Gerard, V. A., DuBois, K., & Greene, R. (1987). Growth responses of two Laminaria saccharina populations to environmental variation. Hydrobiologia, 151(1), 229–232. 10.1007/BF00046134

Ghaderiardakani, F., Quartino, M. L., & Wichard, T. (2020). Microbiome-Dependent Adaptation of Seaweeds Under Environmental Stresses: A Perspective. Frontiers in Marine Science, 7, 575228. 10.3389/fmars.2020.575228

Goldsmit, J., Schlegel, R. W., Filbee-Dexter, K., MacGregor, K. A., Johnson, L. E., Mundy, C. J., Savoie, A. M., McKindsey, C. W., Howland, K. L., & Archambault, P. (2021). Kelp in the Eastern Canadian Arctic: Current and Future Predictions of Habitat Suitability and Cover. Frontiers in Marine Science, 18, 742209. 10.3389/fmars.2021.742209

Harley, C. D. G., Anderson, K. M., Demes, K. W., Jorve, J. P., Kordas, R. L., Coyle, T. A., & Graham, M. H. (2012). Effects of climate change on seaweed communities. Journal of Phycology, 48(5), 1064–1078. 10.1111/j.1529-8817.2012.01224.x

Hsieh, T. C., Ma, K. H., & Chao, A. (2022). iNEXT: iNterpolation and EXTrapolation for species diversity (3.0.0) [Computer software]. http://chao.stat.nthu.edu.tw/wordpress/software-download/

Karsten, U. (2007). Research note: Salinity tolerance of Arctic kelp from Spitsbergen. Phycological Research, 55(4), 257–262. 10.1111/j.1440-1835.2007.00468.x

Kassambara, A. (2022a). ggpubr: “ggplot2” Based Publication Ready Plots. https://CRAN.R-project.org/package=ggpubr

Kassambara, A. (2022b). rstatix: Pipe-Friendly Framework for Basic Statistical Tests. https://CRAN.R-project.org/package=rstatix

King, N. G., Moore, P. J., Thorpe, J. M., & Smale, D. A. (2022). Consistency and Variation in the Kelp Microbiota: Patterns of Bacterial Community Structure Across Spatial Scales. Microbial Ecology. 10.1007/s00248-022-02038-0

Kumar, S., Bhavya, P. S., Ramesh, R., Gupta, G. V. M., Chiriboga, F., Singh, A., Karunasagar, I., Rai, A., Rehnstam-Holm, A.-S., Edler, L., & Godhe, A. (2018). Nitrogen uptake potential under different temperature-salinity conditions: Implications for nitrogen cycling under climate change scenarios. Marine Environmental Research, 141, 196–204. 10.1016/j.marenvres.2018.09.001

Lahti, L., & Shetty, S. (2012). Microbiome R package.

Langdon, C., & Atkinson, M. J. (2005). Effect of elevated pCO2 on photosynthesis and calcification of corals and interactions with seasonal change in temperature/irradiance and nutrient enrichment. Journal of Geophysical Research: Oceans, 110(C9). 10.1029/2004JC002576

Lemay, M. A., Davis, K. M., Martone, P. T., & Parfrey, L. W. (2021). Kelp associated Microbiota are Structured by Host Anatomy. Journal of Phycology, 57(4), 1119–1130. 10.1111/jpy.13169

Lesser, M. P., Fiore, C., Slattery, M., & Zaneveld, J. (2016). Climate change stressors destabilize the microbiome of the Caribbean barrel sponge, Xestospongia muta. Journal of Experimental Marine Biology and Ecology, 475, 11–18. 10.1016/j.jembe.2015.11.004

Li, J., Bates, K. A., Hoang, K. L., Hector, T. E., Knowles, S. C. L., & King, K. C. (2023). Experimental temperatures shape host microbiome diversity and composition. Global Change Biology, 29(1), 41–56. 10.1111/gcb.16429

Lind, A. C., & Konar, B. (2017). Effects of abiotic stressors on kelp early life-history stages. ALGAE, 32(3), 223–233. 10.4490/algae.2017.32.8.7

Lozupone, C. A., & Knight, R. (2007). Global patterns in bacterial diversity. Proceedings of the National Academy of Sciences, 104(27), 11436–11440. 10.1073/pnas.0611525104

Madkaiker, K., Valsala, V., Sreeush, M. G., Mallissery, A., Chakraborty, K., & Deshpande, A. (2023). Understanding the Seasonality, Trends, and Controlling Factors of Indian Ocean Acidification Over Distinctive Bio Provinces. Journal of Geophysical Research: Biogeosciences, 128(1), e2022JG006926. 10.1029/2022JG006926

Mansilla, A., Rosenfeld, S., Rendoll, J., Murcia, S., Werlinger, C., Yokoya, N. S., & Terrados, J. (2014). Tolerance response of Lessonia flavicans from the sub-Antarctic ecoregion of Magallanes under controlled environmental conditions. Journal of Applied Phycology, 26(5), 1971–1977. 10.1007/s10811-014-0294-6

Marshall, K., Joint, I., Callow, M. E., & Callow, J. A. (2006). Effect of Marine Bacterial Isolates on the Growth and Morphology of Axenic Plantlets of the Green Alga Ulva linza. Microbial Ecology, 52(2), 302–310. 10.1007/s00248-006-9060-x

McDevitt-Irwin, J. M., Baum, J. K., Garren, M., & Vega Thurber, R. L. (2017). Responses of Coral-Associated Bacterial Communities to Local and Global Stressors. Frontiers in Marine Science, 4, 262. 10.3389/fmars.2017.00262

McLaren, M. R. (2020). Silva SSU taxonomic training data formatted for DADA2 (Silva version 138) [Dataset]. Zenodo. 10.5281/zenodo.3986799

McLaren, M. R., & Callahan, B. J. (2021). Silva 138.1 prokaryotic SSU taxonomic training data formatted for DADA2 [Dataset]. Zenodo. 10.5281/zenodo.4587955

McMurdie, P. J., & Holmes, S. (2013). phyloseq: An R package for reproducible interactive analysis and graphics of microbiome census data. PLoS ONE, 8(4), e61217.

Mikryukov, V. (2022). metagMisc: Miscellaneous functions for metagenomic analysis (0.0.4) [Computer software].

Monteiro, C. M., Li, H., Bischof, K., Bartsch, I., Valentin, K. U., Corre, E., Collén, J., Harms, L., Glöckner, G., & Heinrich, S. (2019). Is geographical variation driving the transcriptomic responses to multiple stressors in the kelp Saccharina latissima? BMC Plant Biology, 19(1), 513. 10.1186/s12870-019-2124-0

Oksanen, J., Simpson, G. L., Blanchet, F. G., Kindt, R., Legendre, P., Minchin, P. R., O’Hara, R. B., Solymos, P., Stevens, M. H. H., Szoecs, E., Wagner, H., Barbour, M., Bedward, M., Bolker, B., Borcard, D., Carvalho, G., Chirico, M., Caceres, M. D., Durand, S., … Weedon, J. (2022). vegan: Community Ecology Package. https://CRAN.R-project.org/package=vegan

Parada, A. E., Needham, D. M., & Fuhrman, J. A. (2016). Every base matters: Assessing small subunit rRNA primers for marine microbiomes with mock communities, time series and global field samples. Environmental Microbiology, 18(5), 1403–1414. 10.1111/1462-2920.13023

Peteiro, C., & Sánchez, N. (2012). Comparing salinity tolerance in early stages of the sporophytes of a non-indigenous kelp (Undaria pinnatifida) and a native kelp (Saccharina latissima). Russian Journal of Marine Biology, 38(2), 197–200. 10.1134/S1063074012020095

Pita, L., Rix, L., Slaby, B. M., Franke, A., & Hentschel, U. (2018). The sponge holobiont in a changing ocean: From microbes to ecosystems. Microbiome, 6(1), 46. 10.1186/s40168-018-0428-1

Posit team. (2022). *RStudio: Integrated Development Environment for R* (2022.12.0+353) [Computer software]. Posit Software. http://www.posit.co/

Provasoli, L., & Pintner, I. J. (1980). Bacteria Induced Polymorphism in an Axenic Laboratory Strain of Ulva Lactuca (chlorophyceae)1. Journal of Phycology, 16(2), 196–201. 10.1111/j.1529-8817.1980.tb03019.x

Qiu, Z., Coleman, M. A., Provost, E., Campbell, A. H., Kelaher, B. P., Dalton, S. J., Thomas, T., Steinberg, P. D., & Marzinelli, E. M. (2019). Future climate change is predicted to affect the microbiome and condition of habitat-forming kelp. Proceedings of the Royal Society B: Biological Sciences, 286(1896), 20181887. 10.1098/rspb.2018.1887

Quince, C., Lanzen, A., Davenport, R. J., & Turnbaugh, P. J. (2011). Removing Noise From Pyrosequenced Amplicons. BMC Bioinformatics, 12(1), Article 1. 10.1186/1471-2105-12-38

R Core Team. (2022). R: A language and environment for statistical computing [Computer software]. R Foundation for Statistical Computing. https://www.R-project.org/.

Ryan, S. A., Wohlgeschaffen, G. D., Jahan, N., Niu, H., Ortmann, A. C., Brown, T. N., King, T. L., & Clyburne, J. (2019). State of knowledge on fate and behaviour of ship-source petroleum product spills. Fisheries and Oceans Canada = Pêches et océans Canada.

Saha, M., Ferguson, R. M. W., Dove, S., Künzel, S., Meichssner, R., Neulinger, S. C., Petersen, F. O., & Weinberger, F. (2020). Salinity and Time Can Alter Epibacterial Communities of an Invasive Seaweed. Frontiers in Microbiology, 10. https://www.frontiersin.org/articles/10.3389/fmicb.2019.02870

Santoro, E. P., Borges, R. M., Espinoza, J. L., Freire, M., Messias, C. S. M. A., Villela, H. D. M., Pereira, L. M., Vilela, C. L. S., Rosado, J. G., Cardoso, P. M., Rosado, P. M., Assis, J. M., Duarte, G. A. S., Perna, G., Rosado, A. S., Macrae, A., Dupont, C. L., Nelson, K. E., Sweet, M. J., … Peixoto, R. S. (2021). Coral microbiome manipulation elicits metabolic and genetic restructuring to mitigate heat stress and evade mortality. Science Advances, 7(33), eabg3088. 10.1126/sciadv.abg3088

Sogn Andersen, G., Steen, H., Christie, H., Fredriksen, S., & Moy, F. E. (2011). Seasonal Patterns of Sporophyte Growth, Fertility, Fouling, and Mortality of *Saccharina latissima* in Skagerrak, Norway: Implications for Forest Recovery. Journal of Marine Biology, 2011, 1–8. 10.1155/2011/690375

Steneck, R. S., Graham, M. H., Bourque, B. J., Corbett, D., Erlandson, J. M., Estes, J. A., & Tegner, M. J. (2002). Kelp forest ecosystems: Biodiversity, stability, resilience and future. Environmental Conservation, 29(4), 436–459. 10.1017/S0376892902000322

Stratil, S. B., Neulinger, S. C., Knecht, H., Friedrichs, A. K., & Wahl, M. (2014). Salinity affects compositional traits of epibacterial communities on the brown macroalga *Fucus vesiculosus*. FEMS Microbiology Ecology, 88(2), 272–279. 10.1111/1574-6941.12292

Trevathan-Tackett, S. M., Sherman, C. D. H., Huggett, M. J., Campbell, A. H., Laverock, B., Hurtado-McCormick, V., Seymour, J. R., Firl, A., Messer, L. F., Ainsworth, T. D., Negandhi, K. L., Daffonchio, D., Egan, S., Engelen, A. H., Fusi, M., Thomas, T., Vann, L., Hernandez-Agreda, A., Gan, H. M., … Macreadie, P. I. (2019). A horizon scan of priorities for coastal marine microbiome research. Nature Ecology & Evolution, 3(11), 1509–1520. 10.1038/s41559-019-0999-7

Vadillo Gonzalez, S., Hurd, C. L., Britton, D., Bennett, E., Steinberg, P. D., & Marzinelli, E. M. (2024). Effects of temperature and microbial disruption on juvenile kelp Ecklonia radiata and its associated bacterial community. Frontiers in Marine Science, 10. 10.3389/fmars.2023.1332501

van der Loos, L. M., D’hondt, S., Engelen, A. H., Pavia, H., Toth, G. B., Willems, A., Weinberger, F., De Clerck, O., & Steinhagen, S. (2023). Salinity and host drive *Ulva* associated bacterial communities across the Atlantic—Baltic Sea gradient. Molecular Ecology, mec.16462. 10.1111/mec.16462

van Ginneken, V. (2018). Some Mechanism Seaweeds Employ to Cope with Salinity Stress in the Harsh Euhaline Oceanic Environment. American Journal of Plant Sciences, 09(06), 1191–1211. 10.4236/ajps.2018.96089

Weigel, B. L., & Pfister, C. A. (2019). Successional Dynamics and Seascape-Level Patterns of Microbial Communities on the Canopy-Forming Kelp Nereocystis luetkeana and Macrocystis pyrifera. Frontiers in Microbiology, 10. https://www.frontiersin.org/articles/10.3389/fmicb.2019.00346

Weinbauer, M., & Rassoulzadegan, F. (2007). REVIEW: Extinction of microbes: evidence and potential consequences. Endangered Species Research, 3, 205–215. 10.3354/esr003205

Wickham, H. (2016). ggplot2: Elegant Graphics for Data Analysis. Springer-Verlag New York. https://ggplot2.tidyverse.org

Zaneveld, J. R., McMinds, R., & Vega Thurber, R. (2017). Stress and stability: Applying the Anna Karenina principle to animal microbiomes. Nature Microbiology, 2(9), Article 9. 10.1038/nmicrobiol.2017.121

Ziegler, M., Seneca, F. O., Yum, L. K., Palumbi, S. R., & Voolstra, C. R. (2017). Bacterial community dynamics are linked to patterns of coral heat tolerance. Nature Communications, 8(1), Article 1. 10.1038/ncomms14213

